# Interspecific plant competition mediates the metabolic and ecological signature of a plant-herbivore interaction under warming and elevated CO2

**DOI:** 10.1101/420901

**Authors:** Helena Van De Velde, Hamada AbdElgawad, Han Asard, Gerrit T. S. Beemster, Samy Selim, Ivan Nijs, Dries Bonte

## Abstract

1. Biotic interactions shape community evolution, but we lack mechanistic insights on how metabolic and ecological processes under climate change are altered by biotic interactions.
2. We used a two-trophic model community consisting of the aphid *Dysaphis plantaginea* feeding on the forb *Plantago lanceolata*, and a grass competitor *Lolium perenne* that does not experience herbivory by the aphid. Monocultures and mixtures were exposed to the herbivory treatment and to three relevant simulated environmental changes as prevalent under current climate change (increased temperature, CO_2_, and increased temperature and CO_2_)
3. Elevated CO_2_ reduced the nitrogen content of *P. lanceolata*, while simultaneous increases of CO_2_ and temperature modified the plant metabolic component and the magnitude of these responses in different directions. Elevated CO_2_ enhanced defence systems in *P. lanceolata*, but these effects were not altered by warming. Interspecific plant competition did, however, neutralise these responses. There were no indirect effects of climate change on aphid population growth despite changes in plant defense, nutritional quality and biomass induced by our environmental change scenarios.
4. We thus demonstrate interactions between abiotic and biotic processes on plant metabolite profiles, but more importantly, that climate change effect on a selection of the metabolic pathways are altered by herbivory and competition. Our experiment under semi-natural conditions thus demonstrates the non-additive and often neutralizing effects of biotic interactions on plant metabolism and species performance under climate-associated environmental change.

## Introduction

Interactions between insect herbivores and host plants are influenced by primary and secondary metabolites. The concentrations and types of these molecules in plants is likely affected by climate change (Bidart-Bouzat & Imeh-Nathaniel, 2008; Zvereva & Kozlov, 2006). For example, plants exposed to elevated CO_2_ show higher concentrations of many organic molecules, including carbohydrates (fertilization effect), but generally lower or unaltered nitrogen levels, and consequently higher C:N ratios (Bezemer & Jones, 1998; Lincoln *et al*., 1993; Robinson *et al*., 2012; Stiling & Cornelissen, 2007). Lower nitrogen concentrations imply lower levels of leaf protein and amino acids and reduced nutritive value for herbivores (Lincoln *et al*., 1986). Consequently, the performance of insects generally declines under high CO_2_ (Bezemer & Jones, 1998; Hunter, 2001; Lincoln *et al*., 1993). Contrary to elevated CO_2_, moderate warming has positive effects on insect herbivore performance by reducing development time (Bale *et al*., 2002; Van De Velde *et al*., 2016) and increasing fecundity (Meisner *et al*., 2014). Climate change may thus paradoxically constrain rather than facilitate insect herbivore population growth by lowering plant quality (Bauerfeind & Fischer, 2013; Jamieson *et al*., 2015). These resource-quality mediated effects are, however, anticipated to not fully counterbalance the direct effects of warming on development, rendering the overall herbivore response to warming positive (Bauerfeind & Fischer, 2013; Zvereva & Kozlov, 2006).

Secondary metabolites (e.g. lignin, tannins, phenolics and terpenoids), greatly affect tissue quality by determining the nutritive value, palatability, digestibility and/or toxicity of foliage (see Table 1 for a synthesis). Compared to primary metabolites, responses of secondary metabolites to elevated CO_2_ and warming are more variable and less understood (Bidart-Bouzat & Imeh-Nathaniel, 2008; Robinson *et al*., 2012). According to the carbon-nutrient balance hypothesis (Bryant *et al*., 1983), elevated CO_2_ is expected to increase secondary metabolite levels as a result of the ‘excess’ carbon (fertilizing). Yet, because of the relatively lower N-availability, the increase of nitrogen-containing metabolites will be lower than expected (Robinson *et al*., 2012). On the other hand, trade-offs between growth and secondary metabolisms are anticipated to promote primary metabolic activity under warming and therefore favour photosynthesis and growth relative to defence. This expectation corresponds to the growth-differentiation balance hypothesis (Herms & Mattson, 1992). The different components associated with climate change thus induce direct, and often orthogonal responses in chemical defences (Bidart-Bouzat & Imeh-Nathaniel, 2008; Zavala *et al*., 2013; Zvereva & Kozlov, 2006).

**Table 1:** Overview of molecular markers determined in this study, ordered according to their principal function.

| METABOLITE | PRINCIPAL FUNCTION | FUNCTION DESCRIPTION | REF. |
| --- | --- | --- | --- |
| **MARKERS OF OXIDATIVE CELLULAR DAMAGE UNDER ABIOTIC STRESS AND HERBIVORY** |  |  |  |
| **TOCOPHEROL (VITAMINE E)** | Lipophilic antioxidant | Components of biological membranes; act as scavengers of oxygen radicals generated by stress, in particular singlet oxygen. Tocopherols also stabilize membrane structures. | Blokhina et al (2003) |
| **CAROTENOIDS** | Lipophilic antioxidant | Lipophilic antioxidants localized in plastids of photosynthetic and non-photosynthetic plant tissues. Carotenoids react with lipid peroxidation products to end the chain reactions and dissipate the excess excitation energy. | Bartley and Scolnik (1995) |
| **PHENOLICS/ FLAVONOIDS** | Antioxidant | Polyphenols have free radical scavenging activity and are involved in hydrogen peroxide neutralization. Flavonoids are also secondary ROS scavenging molecules in plants under excess excitation. | Blokhina et al (2003) |
| **PROLINE** | Osmolytes/ antioxidant | Proline is an osmolyte, contributing to water uptake, as well as an antioxidant. | Szabados and Savoure (2010). |
| **MALONDIALDEHYDE, (MDA)** | Lipid peroxidation marker | Oxidative stress induces lipid peroxidation disrupting membrane functions. Lipid peroxidation products (MDA) indicate membrane oxidation levels. | Das and Roychoudhury (2014) |
| **TOTAL ANTIOXIDANT CAPACITY (TAC)** |  | Total antioxidant capacity (TAC) is the measure of the gross capacity of a plant extract to scavenge reactive oxygen species. | Das and Roychoudhury (2014) |
| **MARKERS OF HERBIVORY STRESS** |  |  |  |
| **GLYCOSIDES: CATAPOL, AUCUBIN** | Biotic stress defence | Secondary metabolites that help repelling herbivores, protects plant from UV and contributes to control of competitive interactions between plants. | Bowers (1991). |
| **TANNINS** | Biotic stress defence | Tannins are astringent bitter polyphenols and act as feeding deterrents to many insect pests. | Barbehenn et al., (2011) |
| **LIGNIN** | Biotic stress defence | A complex phenolic polymer that plant cell wall rigidity and hydrophobic properties, and is an important barrier against pests and pathogens. | Liu et al., (2018) |
| **SALICYLIC ACID** | Biotic stress defence signal | Salicylic acid is a phytohormone involved in regulation of plant defence, and resistance to insect pests. | War et al., (2012) |
| **JASMONIC ACID** | Biotic stress defence signal | Jasmonic acid is a lipid-derived phytohormone, which protects plants against attacks by a wide variety of herbivores | Zhang et al., 2017 |
| **MARKERS OF WATER STRESS** |  |  |  |
| **SOLUBLE SUGARS (GLUCOSE, SUCROSE, FRUCTOSE)** | Osmolytes | Soluble sugars protect plant cells through osmoregulation, improving water uptake and retention, and may impact cellular membranes through interaction with the lipid bilayer. | Shao et al., (2006) |

Biotic interactions have a strong impact on life histories and physiology of species, and these interactions may shift responses of plants towards climate change (Van de Velde et al. 2016). Competition for limited nutrients for example, enhances the availability of carbon in a focal plant relative to its demand. This, in turn, can increase carbon-based defences and thus reduce herbivory compared with plants that are less constrained by competitors (Bryant *et al*., 1983). Also the identity of a neighbouring plant may affect the resistance of a focal plant by a different outcome of the plant-plant competition (Barton & Bowers, 2006; Broz *et al*., 2010). Conspecific competition, in particular, may result in stronger decline of plant growth and increased defence compared to heterospecific competition. Indeed, Broz *et al*. (2010) for instance showed that focal plants with conspecific neighbours allocated more resources towards production of carbon-based defence molecules, whereas those grown with heterospecific neighbours allocated more resources towards growth. Hence, elevated CO_2_ as well as warming is anticipated to indirectly affect the defence of a focal plant, through species specific stimulation of plant growth and thus plant-plant competition.

Although atmospheric CO_2_ and temperature increase concurrently, empirical studies investigating their combined effect on multi-trophic communities are surprisingly scarce and insufficient to effectively guide theory or synthesis (Cornelissen, 2011; Dyer er al. 2013; Zvereva & Kozlov, 2006). The current study therefore investigates effects of elevated CO_2_ as well as combined effects of elevated CO_2_ and warming on a simple model community consisting of three species: rosy apple aphid *Dysaphis plantaginea* Passerini *(Hemiptera: Aphididae)* feeding on plantain, *Plantago lanceolata* L., and a heterospecific neighboring plant species, perennial ryegrass, *Lolium perenne* L. The aphid does not feed on *L. perenne*. The experimental design consisted of monocultures of *P. lanceolata* and mixtures of *L. perenne* and *P. lanceolata* exposed to the two aforementioned simulated climate scenarios as well as to a control scenario with the current climate. As the experiments were conducted outdoors, we were also able to *post-hoc* study how the imposed climatic and competition treatments affected mildew infections. Biotrophic parasites such as mildew extract nutrients from living cells and have extended periods of physiological interaction with their hosts (Agrios, 2005). Elevated CO_2_ and warming have been shown to affect the severity of pathogen infections (Mikkelsen *et al*., 2015, Thomas & Blanford, 2003). We took advantage of this unplanned infestation to study the impact of elevated CO_2_ and warming on mildew infestation and the putative additive effects of mildew infestation on plant-insect herbivore interaction.

We aim to acquire a more mechanistic understanding of changing ecological interactions under elevated CO_2_ and warming by quantifying a set of relevant metabolites. The metabolite parameters include larger groups (e.g. soluble sugars, tannings, phenolics), as well as more targeted small groups (e.g. tocopherols, carotenoids) and individual molecules (e.g. proline, malondialdehyde). Changes in larger metabolite groups generally indicate more overall metabolic shifts, whereas changes in signature molecules more narrowly indicate particular processes, such as antioxidant defence, abiotic interaction, etc. (see Table 1). Oxidative damage is commonly induced under abiotic stress, but also by herbivory (Wu & Baldwin, 2010). Elevated CO_2_ on the other hand, often reduces this stress impact (Zinta et al. 2018). In more detail, we determined molecular antioxidants (tocopherols, carotenoids, phenolics/flavonoids, proline) and total antioxidant capacity as markers for protection against oxidative stress (Gómez-Ariza *et al*., 2007, Peshev *et al*., 2013). Herbivory damage also induces the production of secondary metabolites associated with the salicylic acid and jasmonic acid pathway (Kuśnierczyk *et al*., 2011; Morkunas & Gabryś, 2011), and the production of protective metabolites that contribute to defence against insects (glycosides, tannins, lignin). Finally, elevated CO_2_ as well as elevated temperature are known to affect stomatal aperture and therefore water transport. To evaluate the plant response to water stress we quantified osmolytes (proline, soluble sugars). Increases in these molecules reduces the cellular water potential, improving water uptake. Given the anticipated complex interplay between elevated CO_2_, temperature and competition, we performed a multivariate grouping analysis to document concerted responses in the quantified metabolites.

Based on the above, we anticipate that elevated CO_2_ and warming will alter foliar nutrients and defence molecules, that altered host quality and plant resistance will affect insect herbivore performance, and that elevated CO_2_ and warming should indirectly influence host quality and plant resistance via effects on neighbouring plants. We predict investments in defence to be higher under conspecific competition relative to heterospecific competition because of increased resource competition. In addition, increased CO_2_ is anticipated to increase chemical defence by promoting the production of secondary metabolites, while simultaneously increased temperature would in contrast promote plant growth. Depending on the relative rates of these physiological changes, net impact on herbivore populations may either remain constant, decline or increase if defence is respectively proportionally promoted or constrained relative to metabolism.

## Materials and methods

### Experimental set-up (Fig S1.1 in appendix 1 from supporting information)

The study was performed at the Drie Eiken Campus, University of Antwerp, Wilrijk, Belgium (51° 09’ N, 04° 24’E) in 12 sunlit, south-facing, climate-controlled chambers (details in Naudts *et al*. 2014). Three climate scenarios (four chambers per scenario) were simulated in an additive design: (1) current atmospheric CO_2_ concentration and temperature (C); (2) future atmospheric CO_2_ and current temperature (CO_2_); and (3) future atmospheric CO_2_ and temperature (TCO_2_). Climate scenarios with elevated CO_2_ had a target CO_2_ concentration of 620 μmol mol^-1^ and future temperature chambers simulated a continuous 3 °C warming above fluctuating ambient temperatures. Climate manipulations were based on the IPCC-SRES B2-scenario prediction of moderate change for the year 2100 (IPCC, 2001).

The CO_2_ concentration in each chamber was continuously measured and maintained at the target concentration with a CO_2_ control group with an infrared analyser (WMA-4, PPSystems, Hitchin, UK). Air temperature and relative humidity were monitored every 0.5 h with a combined humidity–temperature sensor (Siemens QFA66, Erlangen, Germany), by averaging instantaneous readings in half-hour mean values. During the experiment, the CO_2_ concentration was 382 ± 55 μmol mol^-1^ (SD) in the current climate, while it was 615 ± 70 μmol mol^-1^ (SD) in the climate scenarios with future CO_2_ concentration (CO_2_ and TCO_2_). The monthly average air temperature in the C and CO_2_ chambers was 16.2, 17.2 and 18.7 °C in June, July and August, respectively. TCO_2_ chambers were 2.9 ± 1.0 °C (SD) warmer than current temperature chambers. Average vapour pressure deficit was 0.60 ± 0.34 and 0.64 ± 0.52 kPa (SD) in the climate treatments with ambient and warmed air, respectively. Irrigation mimicked the rainfall outside (Naudts et al. 2014), with monthly totals equalling 64.4, 85.1 and 80.2 mm in June, July and August, respectively. Water freely drained while capillary rise was prevented by a drainage system placed below the chambers. Future climate chambers (TCO_2_) received the same amount of water as current climate chambers, so that any increase in water consumption would result in (aggravated) soil drought.

### Plant and insect communities

We used two common co-occurring grassland species, *L. perenne* and *P. lanceolata. P. lanceolata* is characterized by the presence of the iridoid glycosides aucubin and catalpol. These compounds stimulate feeding and oviposition by specialist insects and can act as defence against generalist herbivores (Bowers & Puttick, 1988; Puttick & Bowers, 1988). Both plant species were sown at the end of March in a non-climate controlled greenhouse with a time lag of one week to prevent size differences at the start of the experiment (Cotrufo & Gorissen, 1997), and were watered twice a week. Four or five-week-old seedlings were transplanted into PVC containers (24 cm inner diameter and 40 cm height), filled with sandy soil (93.9% sand, 4.1% silt, 2.0% clay; pH 7.5; Kjeldahl-N 0.125 g kg^−1^; 2.1% C in humus). Each of the 12 chambers received 20 containers with two different compositions: (1) 10 monocultures of *P. lanceolata*, and (2) 10 mixtures of both plant species in a 50:50 ratio. Each community contained 18 individuals planted in a hexagonal grid with 4.5 cm interspace. Interspecific interactions were maximized by avoiding clumping. All communities were fertilized with 10 g m^−2^ NH_4_NO_3_, 5 g m^−2^ P_2_O_5_, 10 g m^−2^ K_2_O and micro-elements (Fe, Mn, Zn, Cu, B, Mo), given dissolved in water in two equal amounts.

When the plants were three months old, *P. lanceolata* was involuntarily infested with powdery mildew *Podosphaera plantaginis*. *P. plantaginis* is a biotrophic fungal pathogen which means that it feeds on living plant tissue, but does not kill the infected host. The powdery mildew was found in all climate scenarios and plant compositions but not all the containers were equally affected. We took advantage of this unplanned infestation to study the impact of elevated CO_2_ and warming on mildew infestation and the putative additive effects of mildew infestation on plant-insect herbivore interaction. Because these infections occurred post-hoc, we do not have proper controls to quantify mildew effects on metabolite composition.

The rosy apple aphid *D. plantaginea* was used as an insect herbivore. It overwinters as eggs on apple trees, the primary host plant, and migrates in spring to the obligate alternate hosts, *P. major* L. and *P. lanceolata* (Alford, 2014). On *Plantago* spp., they give birth to apterous (wingless) morphs that reproduce by parthenogenesis (Blommers *et al*., 2004). *L. perenne* is not a host plant for *D. plantaginea*. The aphids were reared in small cages on *P. lanceolata* under laboratory conditions of 22 ± 1 °C. They were introduced on *P. lanceolata* when the plants were 20 weeks old. At this time, two adult, apterous aphids were placed with a dry paintbrush on the apex of each *P. lanceolata* plant in monocultures and mixtures. Consequently, at the start of the infestation each container was inoculated with 36 (monocultures) or 18 (mixtures) aphids. In each chamber, four monocultures of *P. lanceolata* and four mixtures were randomly chosen for aphid infestation All containers were meshed, and those not receiving aphids acted as controls. The meshes consisted of an 85-cm-tall cylinder of lightweight netting to ensure aphids did not migrate between pots. The infrastructure did not physically limit plant growth and did not cause photosynthetic stress effects (F_v_/F_m_, the intrinsic efficiency of PSII in controls equalled 0.84 which is an optimal value (Johnson *et al*., 1993).

### Data collection

In the fourth week after the aphid introduction we determined the mildew infestation. The degree of powdery mildew on *P. lanceolata* was categorized by a rating system: 1) healthy (no visible lesions); 2) 1% - 25% of the leaves damaged; 3) 26% - 50% of the leaves damaged; 4) 51% - 75% of the leaves damaged; 5) more than 75% of the leaves damaged. At the same time, aphid populations of four replicate communities per plant composition in each chamber were collected (totalling 4 replicate communities × 2 plant compositions × 12 chambers). Aphids were brushed directly into 70% ethanol with a dry paintbrush. It was not possible to collect all the aphids because their number per container was either too high or too low (hard to find), therefore a subsample of the population per container was collected: we searched for and collected aphids for 30 minutes. The total number of aphids per community was divided by the number of *P. lanceolata* individuals in that community. At the same time, we also harvested the plants of two replicate communities per treatment in each chamber (totalling 2 replicate communities × 4 treatments × 12 chambers). Alive aboveground plant biomass in each community was separated from dead biomass by species, dried at 70 °C for 48 h and weighed. These biomass values per species were likewise divided by the number of the species’ individuals in the community, providing primary data for the statistical analysis. The alive biomass samples of *P. lanceolata* plants of the two harvested communities per treatment in each chamber (2 replicate communities × 4 treatments × 12 chambers) were ground in a mill. Three subsamples per community were analysed for nitrogen and carbon content using a NC element analyser (NC-2100 element analyser, Carlo Erba Instruments, Milano, Italy), which were averaged prior to data analysis.

A separate subsample of the milled live aboveground biomass of *P. lanceolata* of the two harvested communities per treatment in each chamber (2 replicate communities × 4 treatments × 12 chambers) was taken to quantify biochemical parameters. These parameters are indicators of stress in the organism (membrane damage as malondialdehyde levels, MDA), indicators of antioxidant defences i.e. total antioxidant capacity (TAC) and antioxidant molecules (tocopherols, carotenes, proline, phenols); indicators of biotic stress and defence, i.e. glycosides (catapol, aucubin), tannin, lignin and salicylic acid; and metabolites involved in plant water-deficit defence (proline, soluble sugars). We refer to Appendix 2 in supporting information for a detailed description of the methods used to quantify these metabolites.

### Data analysis

We focused on *P. lanceolata* because this is the host plant for the aphid and powdery mildew. In a first step we examined covariation in the metabolites responses to the three treatments (aphid infestation, climate scenario and plant composition) by using a hierarchical clustering analysis. Metabolites were normalized by Z-transformation and subjected to a hierarchical clustering analysis with an Euclidean distance metric, and visualized as a heat map using Multi Experiment Viewer (Mev) version 4.8 (Saeed *et al*., 2003). A distance cut off of 0.3 was applied. The clusters were used in the structural equation models (see below).

In a second step, we fitted two piecewise Structural Equation Models (SEM), which combine information from multiple separate linear models into a single causal network (Shipley, 2009). The first SEM investigated the effect of aphid infestation, climate scenario, plant composition and mildew infestation on the metabolites and alive aboveground biomass of *P. lanceolata*. In a second SEM, we separated the metabolites of *P. lanceolata* in control pots from those with aphids. This allowed us to test the effects of the metabolites on the aphid population, as well as to quantify effects of the aphid population on these metabolites. The response of the aphid population was measured as their number at the end of the infestation. The three environmental scenarios are included as a categorical variable and their separate effects on plant performance were tested. Statistics are provided in Appendix 2, and their effects are visualised by coloured arrows in Figure 2. The four clusters obtained by the clustering analysis were used to divide the metabolites in separate groups and hence to avoid redundancy in the analyses. We standardized the metabolites, the alive aboveground biomass of *P. lanceolata* and the number of aphids by converting to Z-scores to equalize variances. To reduce the number of mildew categories, we rearranged degree of mildew in two categories: (1) no or mild mildew infection (category 1 - 2) (2) severe mildew infection (category 3 - 5).

Traditional SEM estimation methods assume that all variables follow a normal distribution and all observations are independent (Grace, 2006). In our analyses, we used piecewise SEM that allows fitting general linear mixed effect models that can incorporate random effects. Each mixed effects model was fitted using the “lme” function in the “nlme” package (version 3.1-128) in R. As outline higher, multiple containers from each of the four competition x herbivory treatments were located in a single chamber. To avoid pseudoreplication, chamber was been added as a random effect in all models. The overall path model (the SEM) was fitted using the “piecewiseSEM” package (version 1.2.1) in R (Lefcheck, 2016). Goodness of fit was estimated using Shipley’s test of d-separation, which yields a Fisher’s C statistic that is Chi-squared distributed (Shipley, 2009). If the resulting P-value > 0.05, then the SEM can be said to adequately reproduce the hypothesized causal network.

As a final confirmatory step, all data were analysed with General Linear Mixed models (GLM) in SAS (version 9.2, SAS Institute Inc., Cary, NC) (Littell *et al*., 1996) with chamber as a random factor nested within climate scenario. Climate scenario, plant composition, aphid infestation and two-way and three-way interactions between these predictors were included as fixed factors. Because mildew infestation did not have a significant effect on biochemical plant responses, this treatment was excluded from the GLM analysis. Non-significant factors were backwards-excluded from the model. In case of significant effects, *a posteriori* means comparisons using the Tukey test, corrected for multiple comparisons, were made. Effects were considered significant at P ≤ 0.05. Repeated univariate analyses are subject to error proliferations. We deliberately decided not to correct results from these analyses for multiple testing, but instead interpret them in a conservative way in congruence with the higher described grouping analyses. Results and discussion of the series of univariate analyses are therefore provided in Appendix 3 from supporting information.

## Results

### Hierarchical clustering analysis of metabolites

The variation in primary and secondary metabolites of *P. lanceolata* was subjected to a hierarchical clustering analysis in order to understand their grouping according to their covariation in response to the imposed herbivory, climate and competition treatments. We retained both specific soluble sugars as well as their grouped responses as both may differently respond among treatments. We obtained four major clusters of metabolites (clusters 1 - 4 in Fig. 1), which were not affected by leaving out the total amount of soluble sugars (see Fig S1.2 in supporting information).

**Fig. 1.**
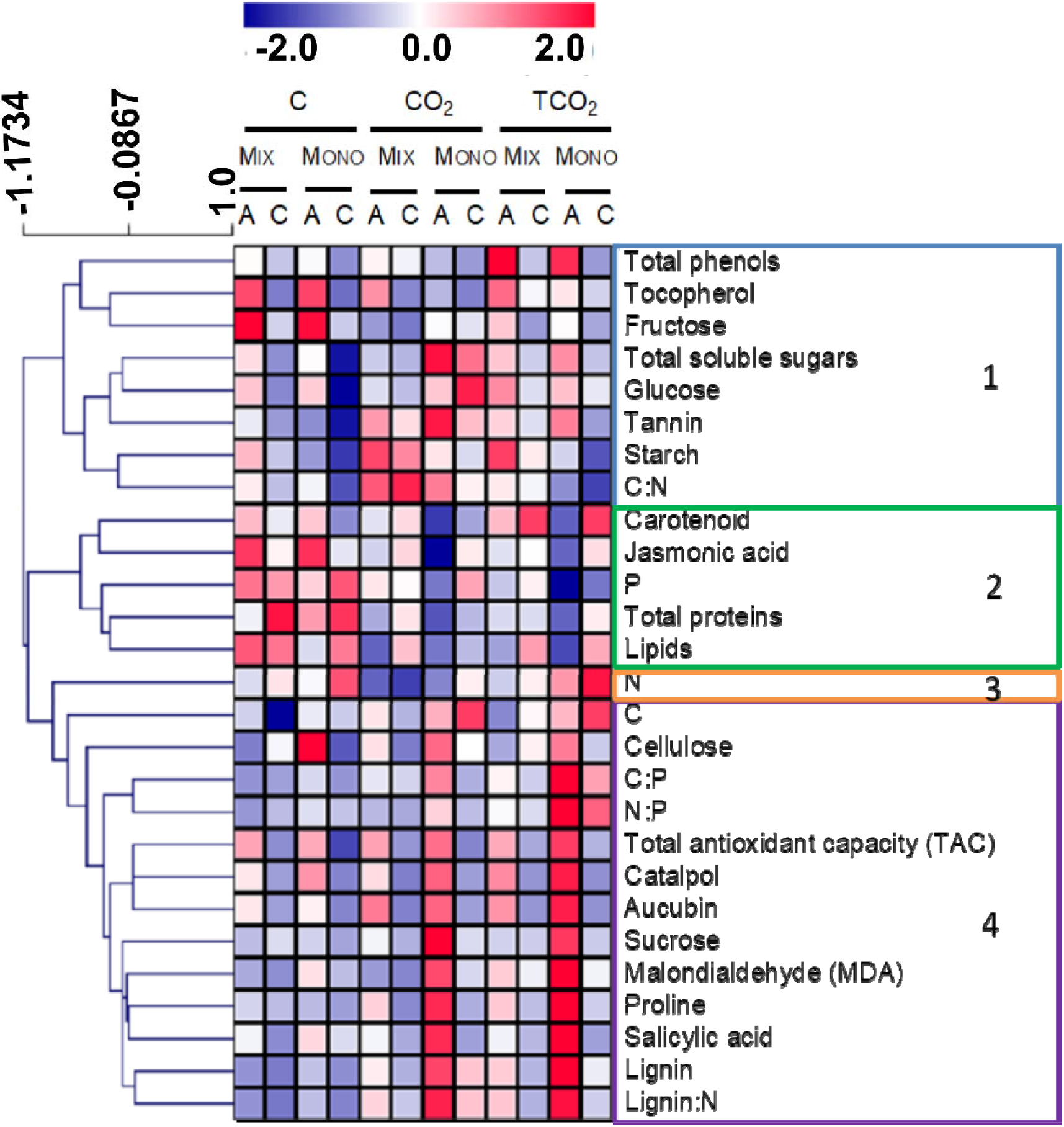
Heat map showing the metabolite levels in the leaves of P. lanceolata, normalized to Z-score for each metabolite (blue-white-red heat map). Blue and red colours indicate a low and high metabolite level, respectively. Clustering was based on the Euclidean distance for metabolites. Labels 1-4 and colours indicate the four prominent clusters. Labels C, CO_2_ and TCO_2_ indicate current climate, elevated CO_2_ and combined warming and elevated CO_2_, respectively. Plant communities consist of monocultures of P. lanceolata (mono) and mixtures of Lolium perenne and P. lanceolata (mix) with (A) and without aphids (C).

The first cluster containing metabolites consistently upregulated by the presence of aphids contains the phenol antioxidants, the lipophilic antioxidant tocopherol, the C:N ratio, tannins, and the non-structural carbohydrates soluble sugars (including glucose, fructose) and starch (Fig. 1). The second cluster contains another group of antioxidant and defence molecules that are downregulated by elevated CO_2_ and more so by aphids in mono-cultures, i.e. the carotenoids, as well as the growth regulator, jasmonic acid, total proteins, phosphorous and lipids (Fig. 1).

The nutrient N which contributes solely to the third cluster, is downregulated under elevated CO_2_ . Carbon concentration, the C:P and N:P ratio, and lignin:N were classified in the fourth cluster that is specifically upregulated by aphids in monocultures under elevated CO_2_ (Fig. 1). Cluster 4 also contains metabolites that play a role in plant defences against herbivores, such as salicylic acid and the iridoid glycosides (catalpol and aucubin), and also contains TAC, lignin, cellulose, proline and MDA. Notably in this cluster, metabolites are strongly induced in CO_2_ and TCO_2_.

### Causal changes as inferred by structural equation models and univariate analyses

SEMs quantify the strength of both the direct and indirect interactions between the environmental scenarios related to climate change, herbivory and plant phenotype. The first SEM presents the effect of aphid infestation (qualitative effect), climate scenario, plant composition and mildew infestation on the chemical composition (the metabolites of *P. lanceolata* were subdivided into the four groups obtained from the hierarchical clustering; see Fig. 1) and the live aboveground biomass of *P. lanceolata* (Fig. 2A). The hypothesized structural relationship with climate scenario affecting mildew infection severity, and climate scenario, plant composition and aphid infestation interactively affecting metabolites and biomass adequately fits the data (χ^2^ = 42.53, df = 32, p = 0.101).

**Fig. 2.**
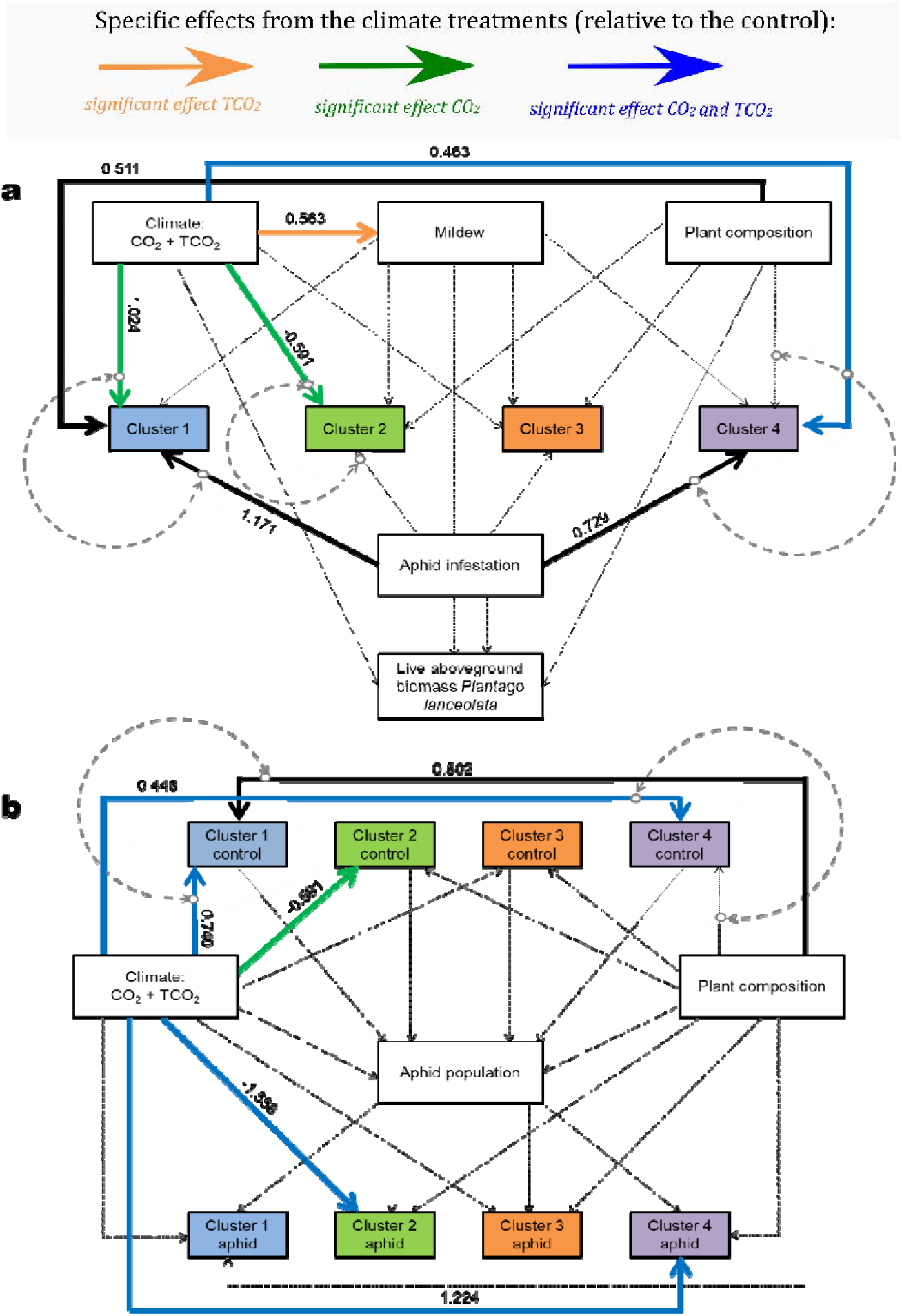
(a) Structural equation model showing how climate scenario (CO_2_ and TCO_2_), mildew infestation, plant composition and aphid infestation affect the chemical composition and the live aboveground biomass of P. lanceolata. (b) Structural equation model showing how climate scenario, plant composition and chemical composition of P. lanceolata affect aphid population and how aphid population, in turn, affects the chemical composition of P. lanceolata. The four clusters refer to those obtained by the hierarchical clustering analysis (see Fig. 1). Solid black, green and orange arrows represent significant relationships (P ≤0.05) and dashed grey lines significant interactions. Blue lines stand for significant effects of both CO_2_ and TCO_2_, green lines for significant effects of CO_2_ and orange lines for significant effects of TCO_2_. Light grey arrows represent nonsignificant relationships. Standardized path coefficients are shown next to pathways. For the effect of CO_2_ and TCO_2_, the average path coefficients are shown. The individual path coefficients of CO_2_ and TCO_2_ can be seen in Table S2.1 and Table S2.2 (see Appendix 2). Metabolites levels, live aboveground biomass of P. lanceolata and number of aphids were scaled before analysis.

Based on this SEM (see Fig. 2A, Table S2.1 in Appendix 2 and the univariate analyses, we draw the following key-insights:

1. Environmental change scenario, nor aphid infestation or plant composition had a significant impact on the live aboveground biomass of *P. lanceolata*. Live biomass tended to be higher in CO_2_ relative to the control C and was significantly lower in TCO_2_ than CO_2_, which combined lead to similar values in TCO_2_ and the control C (Fig. 3; see also Appendix 3). No direct impacts of the environmental change scenarios on aphid population growth were recorded, and neither did elevated CO_2_ and temperature change the impact of herbivory on *P. lanceolata* biomass. Despite absence of any response in the ecological endpoints, we found climate scenario, aphid infestation and plant composition to impact the distinguished clusters in a different manner.
2. An increased CO_2_ not associated with increased temperatures, herbivory and plant competition impact oxidative stress and water stress related metabolites from cluster 1, while the anti-oxidants from cluster 2 were only impacted by the increased CO_2_. We noticed a significant interaction between climate scenario and aphid infestation for metabolites associated with cluster 1 (see Table S3.1 and Fig. S3.1 in appendix 3 from the supporting information). Overall, aphid infestation increased the concentrations of the metabolites pending on the environmental change scenario between 30 and 100% in cluster 1 (except for C:N ratio; Fig. 4A-B). The concentrations of glucose and total soluble sugars increased at elevated CO_2_ but only in monocultures, while TCO_2_ did not alter the concentrations of glucose relative to the CO_2_ treatment but decreased the concentrations of soluble sugars in monocultures (Fig. 4C). In general CO_2_, compared to the control, increased the concentration of tannin while TCO_2_ did not impose further differences. Increased CO_2_ also impacted especially the proteins and lipids from cluster 2. Relative to the control, CO_2_ decreased the concentration of proteins irrespective of the plant composition while aphid infestation and TCO_2_, compared to CO_2_, did not alter it. The concentration of lipids decreased with aphid infestation in monocultures and mixtures in all climate scenarios (except for mixtures in C).
3. Elevated temperature and CO_2_ increased the mildew infestation compared to the control of no environmental change (C), but the mildew infestation, in turn, did not affect the metabolites and the alive aboveground biomass of *P. lanceolata*. Under this environmental change, and in accordance with the hypothesis that metabolites from the second cluster are typically associated with herbivore stress, single responses are conditional to aphid infestation and plant competition as well (see Table S3.2, Fig. S3.2 in supporting information). More specifically, concentrations of jasmonic acid increased with aphid infestation under the control conditions, but its response was depressed under elevated temperature and CO_2_, especially in monocultures (Fig. 4D). Interspecific interactions between *P. lanceolata* and *L. perenne* in consequence mitigated the negative effect of aphid infestation in these climate scenarios, and hence have a neutralizing effect. Aphid infestation reduced the concentration of P in monocultures, but did not alter it in mixtures (Fig. 4E). Furthermore, aphid infestation decreased the concentration of proteins in mixtures in absence of the imposed environmental changes but did not alter the concentrations in monocultures and mixtures in CO_2_ and TCO_2_ (Fig. 4F). While CO_2_, compared to the control C, decreased the concentration of lipids in monocultures without aphids and mixtures with aphids, its concentration increased again to similar levels in TCO_2_, compared to CO_2_ and control monocultures without aphids. Antioxidant Carotenoid concentration increased in TCO_2_, compared to the control C and CO_2_. Elevated temperature and CO_2_ reduced the concentration of starch (except for mixtures with aphids) and the C:N ratio compared to CO_2_.
4. Leaf nitrogen was not impacted by any of the treatments (cluster 3; see Table S3.3 in supporting information).
5. Aphid infestation, and CO_2_ increase, independent of a change in temperature increased the metabolites in cluster 4 metabolites that play a role in plant defences against herbivores and oxidative stress resistance (see Table S3.4, Fig. S3.3 & Fig. S3.4 in supporting information). They jointly inhibited the expression of salicylic acid under the two environmental change scenarios (Fig 4G). Besides these single factor treatments effects, climate scenario, aphid infestation and plant composition also interacted with each other in affecting the metabolites in cluster 4. CO_2_ and TCO_2_ strengthened (salicylic acid and lignin) or induced (proline) an effect of aphid infestation but only in monocultures (Fig. 4G). TCO_2_, compared to CO_2_ and C increased the N:P and the C:P ratio in monocultures (Fig. 4H,I). Cellulose levels were always higher in monocultures and aphid infestation but the magnitude of increase was lowers under the two climate-related environmental change scenarios relative to the current conditions (control).

**Fig. 3.**
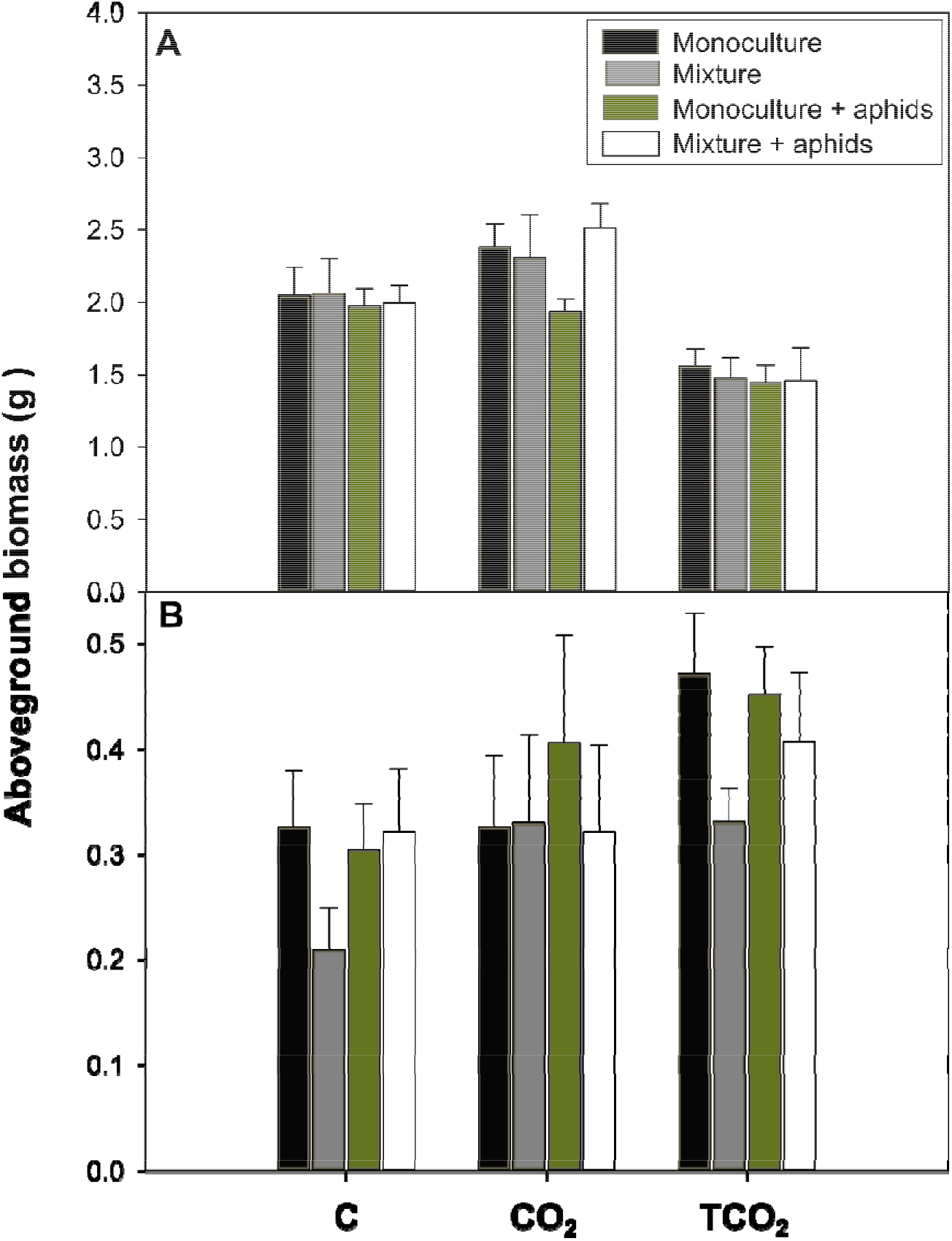
Effect of climate scenario (C, CO_2_ and TCO_2_), aphid infestation and plant composition on the live (A) and dead (B) aboveground biomass of individual P. lanceolata plants. Bars represent means ± SE. Plant communities consist of monocultures of P. lanceolata and mixtures of Lolium perenne and P. lanceolata.

**Fig. 4.**
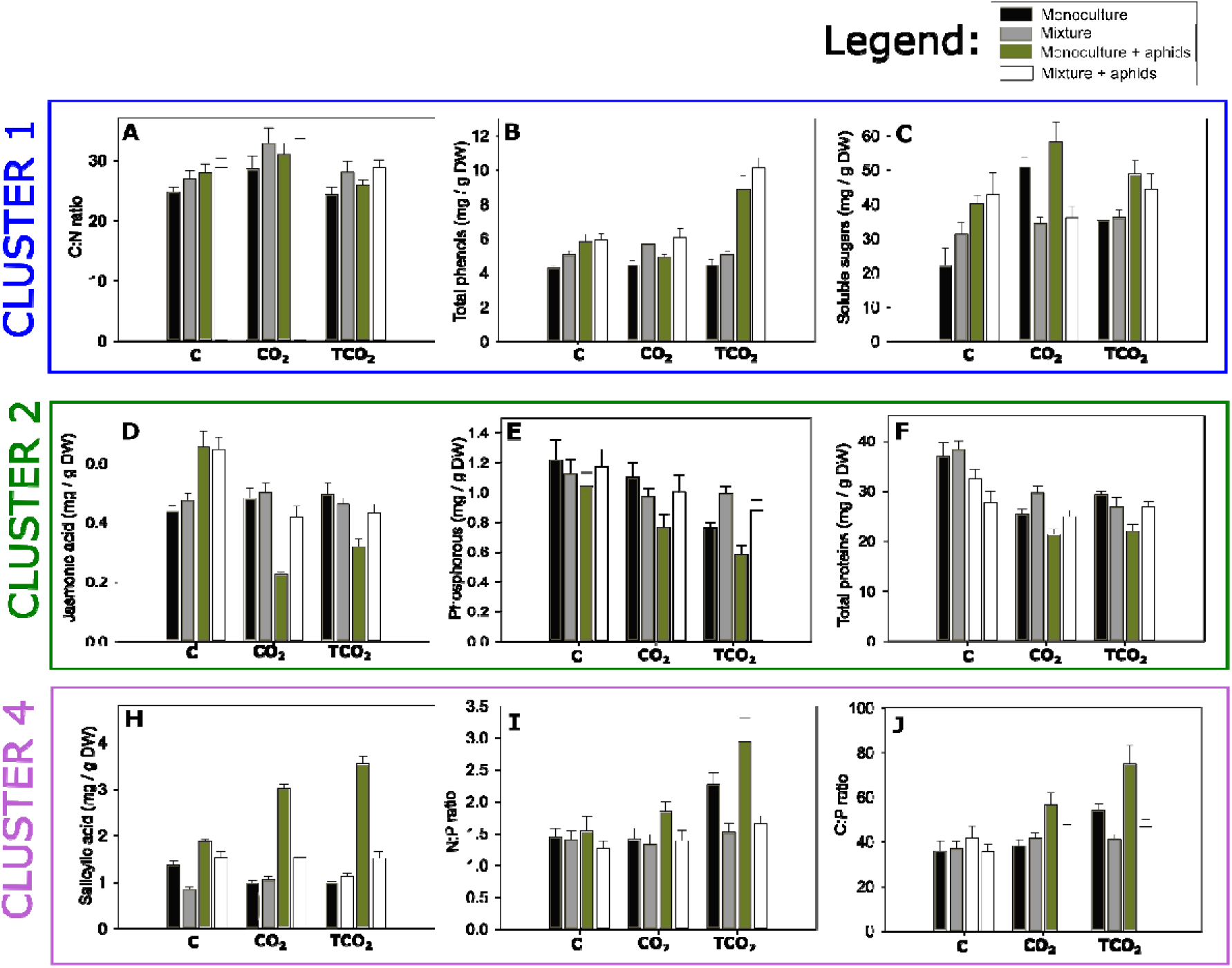
Effect of climate scenario (C, CO_2_ and TCO_2_), aphid infestation and plant composition on a selection of relevant metabolites from the four distinguished clusters. Bars represent means ± SE. Plant communities consist of monocultures of P. lanceolata and mixtures of Lolium perenne and P. lanceolata. As cluster 3 only contained one metabolite, it is not depicted here.

The second SEM presents the effect of climate scenario, plant composition and chemical composition of *P. lanceolata* on the aphid population and whether the aphid population, in turn, altered the chemical composition of *P. lanceolata* (quantitative effect). The mildew infestation was left out of this analysis since it is a parasite of the plants, not of the aphids. The results of this SEM model show the following goodness of fit statistics: χ^2^ = 77.52, df = 80 and p = 0.558 (Table S2.2 and Fig. 2b), which provides us with two insights. Firstly, climate scenario, plant composition and the chemical composition of *P. lanceolata* did not affect the number of aphids. Secondly, the aphid population size in turn, did not change the chemical composition of *P. lanceolata*. Bringing together the findings from both SEM models, we conclude that the treatment aphid infestation altered the chemical composition of *P. lanceolata* but aphid population size had no effect on it.

## Discussion

A future climate clearly altered leaf nutritional quality and the expression of defence molecules, while aphid infestation overall impaired nutritional requirements for insect herbivores. Despite the strong metabolic changes as induced by climate-related environmental change, no feedbacks on aphid population performance and herbivory were detected. More importantly, we show that interspecific plant competition neutralizes the positive effects of elevated CO_2_ on the production of defence molecules in *P. lanceolata*. Induced effects of climate-related environmental change on plant performance can thus be altered and even nullified by competitive and food web interactions. We first discuss the general herbivore-induced responses followed by an in-depth discussion of their dependency on the imposed climate and competition treatments.

### Herbivore-induced responses

Our experiment, as conducted under unique semi-natural conditions confirmed in first instance findings that aphid infestation reduces N, P, total proteins and lipids in *P. lanceolata* leaves, thereby impairing nutritional requirements of insect herbivores (Schoonhoven *et al*., 2005). In contrast to previous research (Walling, 2000), however, aphid infestation promoted the salicylic acid and jasmonic acid pathway (Kuśnierczyk *et al*., 2011; Morkunas & Gabryś, 2011) as well as an increased production of catalpol, aucubin, tannin, lignin and cellulose.

Aphid herbivory increased oxidative stress in plants as well (Wu & Baldwin, 2010). We assessed the induction of oxidative stress by measuring MDA, which aphid herbivory enhanced considerably. A common defence response of plants against oxidative stress includes the increase of antioxidant metabolites. Indeed, TAC and the well-known antioxidant molecules phenols and tocopherols increased due to aphid herbivory. Moreover, *P. lanceolata* responded to herbivory by increasing soluble sugars. At higher concentrations, also sugars can act as antioxidants and may play a signalling role in regulating stress and defence responses (Gómez-Ariza *et al*., 2007, Peshev *et al*., 2013). The induced resistance against oxidative stress associated with aphid feeding appeared sufficiently effective to constrain biomass loss from herbivory.

### Climate dependency of the plant phenotype

Elevated CO_2_ and temperature did not change the impact of herbivory on *P. lanceolata* biomass. This implies that warming and CO_2_ did not affect the net interaction strength between plants and herbivores under semi-natural conditions. In contrast to earlier studies (Van De Velde *et al*., 2016), our experiment allows a profound analysis of the underlying biochemical processes.

First, we found elevated CO_2_ to increase starch and soluble sugars (in monocultures) and to lower leaf proteins, hereby increasing the C:N ratio. In addition, elevated CO_2_ reduced P and consequently increased the C:P ratio (in monocultures), thereby reducing the plant’s nutritive value to herbivores (Huberty & Denno, 2006; Lincoln *et al*., 1986; Mattson, 1980; Schoonhoven *et al*., 2005). We demonstrate here that these effects can be countered when elevated CO_2_ is accompanied with a temperature rise. This contradicts findings based on a limited set of primary metabolites (Murray *et al*. 2013), but urges to apply broader phenomic approaches to fully understand plant responses to climate change.

Second, elevated CO_2_ increased the defence molecules lignin and tannin, especially under aphid infestation. The induction of plant chemical defences (Bidart-Bouzat & Imeh-Nathaniel, 2008; Bidart-Bouzat *et al*., 2005) is thus not consistent for all defence molecules, and are consequently anticipated to render prediction on future plant-insect herbivore interaction highly complex (Agrawal, 1999). In our study, the higher C:N ratio reflected higher lignin and tannin levels. While there is a consensus that the carbon-nutrient-balance hypothesis fails as predictive framework (Hamilton *et al*., 2001; Lindroth, 2012), alternative hypotheses have been proposed stating that resource utilization for chemical defence is linked with photosynthesis, hormone regulation and the control of gene expression (Zavala *et al*., 2017). We here provide evidence for this alternative mechanism as we found CO_2_ to alter phytohormones (i.e., salicylic acid and jasmonic acid) that play an important role in promoting compounds responsible for herbivore defence (Wu & Baldwin, 2010). Since no further quantitative changes were observed under higher temperature, we can confidently conclude that climate-change-induced changes in metabolic profiles are more due to enhanced CO_2_ than to temperature increases acting in parallel.

### Competition: a biotic interaction mediating climate-induced effects

The presence and identity of neighbouring plants can influence the quality of the host plant by altering primary and secondary chemistry (Barton & Bowers, 2006; Broz *et al*., 2010, Lankau, 2012; Thorpe *et al*., 2011). We provide evidence that these biotic interactions are additionally able to alter climate-induced phenomic changes, and by that further impacting ecological interactions under climate-related environmental change. Energetic growth-defence-trade-offs have usually been attributed to these competition effects (Herms & Mattson, 1992), but recent studies demonstrated the role of light limitation for plant defence downregulation (Ballaré, 2014; Campos *et al*., 2016). More specifically, a reduced red to far-red ratio downregulates defences by a simultaneous inhibition of the jasmonic acid and the salicylic acid pathway (Wit *et al*., 2013). The lower levels of defence molecules and salicylic acid in *P. lanceolata* support this hypothesis. Competition did, however, also counter the CO_2_-induced changes in primary and secondary metabolites, as well as those induced by herbivores, rendering an understanding of plant defences far from trivial (de Vries *et al*., 2017). It is well-understood that bottom-up effects from plant quality to herbivore abundance cannot be generalized across the feeding guilds (Bezemer & Jones, 1998; Robinson *et al*., 2012; Stiling & Cornelissen, 2007), but phloem feeders such as aphids were at least expected to be facilitated. Despite the complex, high-dimensional changes in plant metabolomes under competition and climate change, the overall effects on aphid population sizes were small to non-existent.

We here thus demonstrate that the joint action of atmospheric factors associated with climate change and biotic interactions are able to induce pronounced changes in plant metabolomes and biomass, but that translation of these biochemical changes towards ecological responses appears non-trivial. Climate-change-imposed changes are shown to be completely neutralized when common biotic interactions are taken into consideration. Plant-plant interactions thus add a layer of complexity in mechanistic studies of climate change effects on plant-enemy ecological interaction.

## SUPPORTING INFORMATION

Additional supporting information may be found in the online version of this article.:

**Appendix 1:** Schematic representation of the experimental setup and clustering without total amount of soluble sugars

**Appendix 2:** Metabolite quantification methods and SEM statistics

**Appendix 3:** Results from the univariate GLM analyses (statistics and figures)

## Acknowledgements

H. Van De Velde is a Research Assistant of the Fund for Scientific Research-Flanders (FWO). D. Bonte was funded by the FWO research network W0.003.16N “An eco-evolutionary network of biotic interactions”. We thank the Earth and Life Institute (Université Catholique de Louvain) for providing *Dysaphis plantaginea*, N. Calluy and M. Wellens for technical assistance, colleagues for field assistance and Martijn Bezemer and Martijn Vandegehuchte for helpful comments on an earlier version of this manuscript.

