## appendix 3 for "Interspecific plant competition mediates the metabolic and ecological signature of a plant-herbivore interaction under warming and elevated CO2"

**Appendix 3: Results from the univariate GLM analyses**

**3.1 Aboveground biomass:** Climate scenario, but not plant composition or aphid infestation altered the live aboveground biomass of *P. lanceolata* (F_2,9_ = 6.34, P= 0.019; F_1.82_ = 0.84, P = 0.362; F_1.82_ = 0.98, P= 0.325, respectively, Fig. 3A). The live aboveground biomass tended to be higher in CO_2_ than C and was significantly lower in TCO_2_ than CO_2_, which combined led to similar values in TCO_2_ and C. Plant composition, but not climate scenario or aphid infestation altered the dead aboveground biomass of *P. lanceolata* (F_1,75_ = 72.82, P= <0.001; F_2.9_ = 0.93, P= 0.428; F_1,75_ = 0.71, P= 0.401, respectively, Fig. 3B). Competition from *L. perenne* reduced the dead aboveground biomass relative to growing with conspecific neighbors. Furthermore, climate scenario and plant composition did not alter relative herbivory effects (F_2.42_ = 0.02, P= 0.981; F_1.42_ = 2.37, P= 0.131, respectively). In addition, climate scenario and plant composition did not alter the number of aphids per plant (F_2,9_ = 0.45, P= 0.650; F_1,78_ = 0.87, P= 0.353, respectively).

**3.2 Metabolites from the different clusters**

**Table S3.1** Summary of GLM results for effects of climate scenario, plant composition and aphid infestation on the metabolite levels in cluster 1. Plant communities consist of monocultures of *P. lanceolata* and mixtures of *Lolium perenne* and *P. lanceolata*. P-values are presented in bold when significant (≤ 0.05).

| Measurement | Treatment | df | F | P |
| --- | --- | --- | --- | --- |
| Total phenols | Climate scenario | 2,84 | 12.68 | **<0.001** |
|  | Plant composition | 1,84 | 23.24 | **<0.001** |
|  | Aphid infestation | 1,84 | 79.59 | **<0.001** |
|  | Climate scenario × aphid infestation | 2,84 | 18.15 | **<0.001** |
|  | Climate scenario × plant composition | 2,84 | 1.41 | 0.250 |
|  | Plant composition × aphid infestation | 1,84 | 2.40 | 0.125 |
|  | Climate scenario × aphid infestation × plant composition | 2,84 | 0.17 | 0.845 |
| Tocopherols | Climate scenario | 2,84 | 15.84 | **<0.001** |
|  | Plant composition | 1,84 | 18.30 | **<0.001** |
|  | Aphid infestation | 1,84 | 196.73 | **<0.001** |
|  | Climate scenario × aphid infestation | 2,84 | 22.72 | **<0.001** |
|  | Climate scenario × plant composition | 2,84 | 3.56 | **0.003** |
|  | Plant composition × aphid infestation | 1,84 | 6.70 | **0.011** |
|  | Climate scenario × aphid infestation × plant composition | 2,84 | 3.89 | **0.024** |
| Fructose | Climate scenario | 2,84 | 8.10 | **0.001** |
|  | Plant composition | 1,84 | 0.4 | 0.529 |
|  | Aphid infestation | 1,84 | 17.66 | **<0.001** |
|  | Climate scenario × aphid infestation | 2,84 | 4.88 | **0.001** |
|  | Climate scenario × plant composition | 2,84 | 1.14 | 0.323 |
|  | Plant composition × aphid infestation | 1,84 | 0.12 | 0.732 |
|  | Climate scenario × aphid infestation × plant composition | 2,84 | 0.08 | 0.928 |
| Soluble sugars | Climate scenario | 2,84 | 7.57 | **0.001** |
|  | Plant composition | 1,84 | 4.97 | **0.028** |
|  | Aphid infestation | 1,84 | 19.17 | **<0.001** |
|  | Climate scenario × aphid infestation | 2,84 | 1.69 | 0.190 |
|  | Climate scenario × plant composition | 2,84 | 10.68 | **<0.001** |
|  | Plant composition × aphid infestation | 1,84 | 1.59 | 0.210 |
|  | Climate scenario × aphid infestation × plant composition | 2,84 | 0.02 | 0.985 |
| Glucose | Climate scenario | 2,84 | 4.20 | **0.018** |
|  | Plant composition | 1,84 | 0.63 | 0.428 |
|  | Aphid infestation | 1,84 | 8.01 | **0.006** |
|  | Climate scenario × aphid infestation | 2,84 | 6.77 | **0.002** |
|  | Climate scenario × plant composition | 2,84 | 3.81 | **0.013** |
|  | Plant composition × aphid infestation | 1,84 | 0.24 | 0.625 |
|  | Climate scenario × aphid infestation × plant composition | 2,84 | 2.05 | 0.136 |
| Tannin | Climate scenario | 2,84 | 23.51 | **<0.001** |
|  | Plant composition | 1,84 | 0.22 | 0.638 |
|  | Aphid infestation | 1,84 | 15.55 | **<0.001** |
|  | Climate scenario × aphid infestation | 2,84 | 2.07 | 0.132 |
|  | Climate scenario × plant composition | 2,84 | 8.61 | **<0.001** |
|  | Plant composition × aphid infestation | 1,84 | 9.28 | **0.003** |
|  | Climate scenario × aphid infestation × plant composition | 2,84 | 0.06 | 0.941 |
| Starch | Climate scenario | 2,84 | 23.51 | **<0.001** |
|  | Plant composition | 1,84 | 75.83 | **<0.001** |
|  | Aphid infestation | 1,84 | 35.69 | **<0.001** |
|  | Climate scenario × aphid infestation | 2,84 | 2.29 | 0.108 |
|  | Climate scenario × plant composition | 2,84 | 1.08 | 0.344 |
|  | Plant composition × aphid infestation | 1,84 | 0.62 | 0.435 |
|  | Climate scenario × aphid infestation × plant composition | 2,84 | 0.04 | 0.964 |
| C:N ratio | Climate scenario | 2.84 | 8.71 | **<0.001** |
|  | Plant composition | 1,84 | 6.74 | **0.011** |
|  | Aphid infestation | 1,84 | 2.43 | 0.123 |
|  | Climate scenario × aphid infestation | 2,84 | 0.38 | 0.684 |
|  | Climate scenario × plant composition | 2,84 | 0.31 | 0.733 |
|  | Plant composition × aphid infestation | 1,84 | 1.12 | 0.293 |
|  | Climate scenario × aphid infestation × plant composition | 2,84 | 0.17 | 0.842 |

**Fig. S3.1** Effect of climate scenario (C, CO_2_ and TCO_2_), aphid infestation and plant composition on the metabolite levels in cluster 1. Bars represent means ± SE. Plant communities consist of monocultures of *P. lanceolata* and mixtures of *Lolium perenne* and *P. lanceolata*.

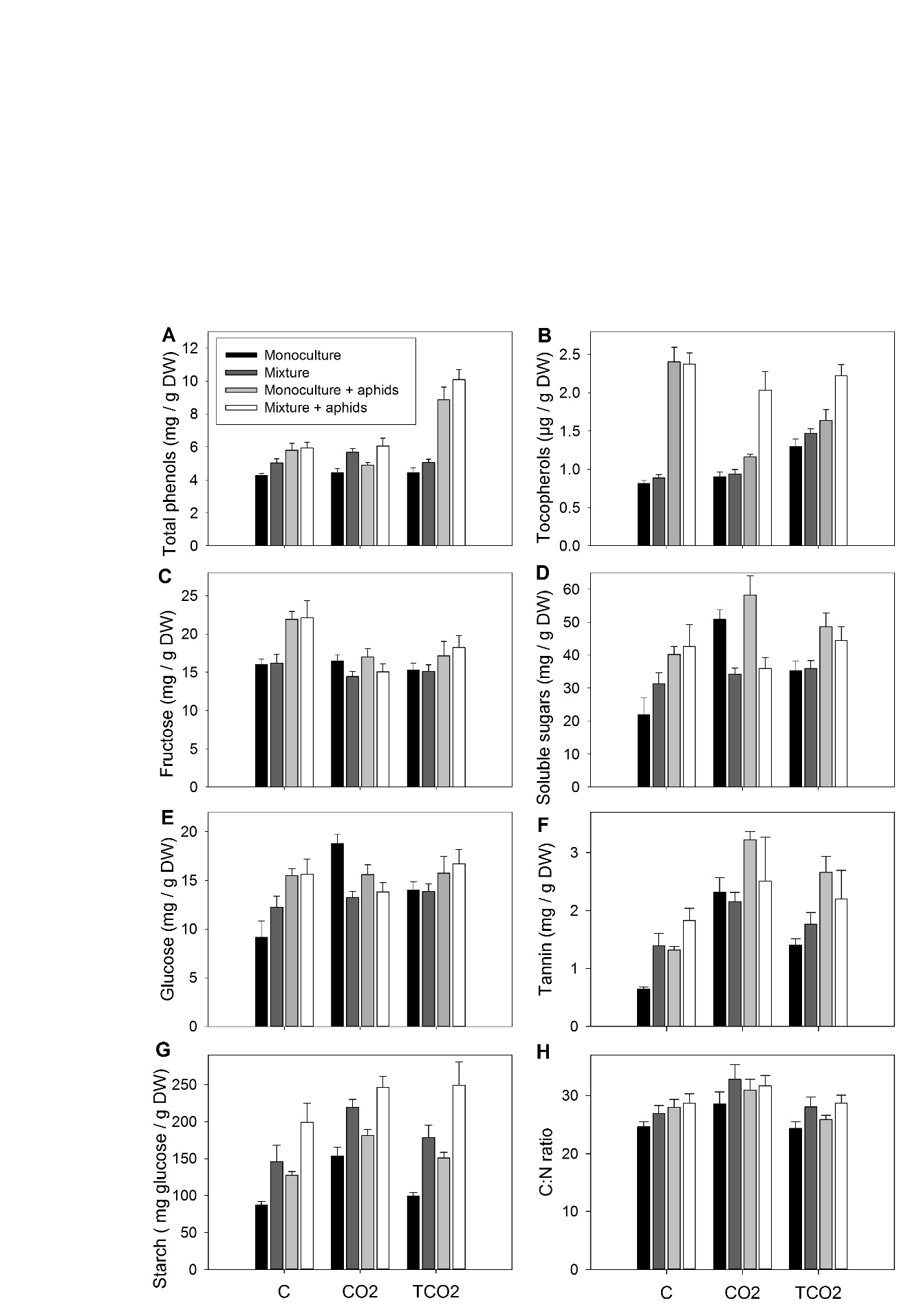

**Table S3.2** Summary of GLM results for effects of climate scenario, plant composition and aphid infestation on the metabolite levels in cluster 2. Plant communities consist of monocultures of *P. lanceolata* and mixtures of *Lolium perenne* and *P. lanceolata*. P-values are presented in bold when significant (≤ 0.05).

| Measurement | Treatment | df | F | P |
| --- | --- | --- | --- | --- |
| Carotenoids | Climate scenario | 2,84 | 9.17 | **<0.001** |
|  | Plant composition | 1,84 | 14.48 | **<0.001** |
|  | Aphid infestation | 1,84 | 5.67 | **0.019** |
|  | Climate scenario × aphid infestation | 2,84 | 15.28 | **<0.001** |
|  | Climate scenario × plant composition | 2,84 | 1.06 | 0.351 |
|  | Plant composition × aphid infestation | 1,84 | 1.30 | 0.257 |
|  | Climate scenario × aphid infestation × plant composition | 2,84 | 2.85 | 0.064 |
| Jasmonic acid | Climate scenario | 2,84 | 24.74 | **<0.001** |
|  | Plant composition | 1,84 | 8.62 | **0.004** |
|  | Aphid infestation | 1,84 | 2.06 | 0.154 |
|  | Climate scenario × aphid infestation | 2,84 | 37.38 | **<0.001** |
|  | Climate scenario × plant composition | 2,84 | 2.18 | 0.120 |
|  | Plant composition × aphid infestation | 1,84 | 6.08 | **0.016** |
|  | Climate scenario × aphid infestation × plant composition | 2,84 | 3.53 | **0.034** |
| Phosphorous | Climate scenario | 2,84 | 14.30 | **<0.001** |
|  | Plant composition | 1,84 | 4.87 | **0.030** |
|  | Aphid infestation | 1,84 | 5.53 | **0.021** |
|  | Climate scenario × aphid infestation | 2,84 | 0.34 | 0.716 |
|  | Climate scenario × plant composition | 2,84 | 2.21 | 0.116 |
|  | Plant composition × aphid infestation | 1.84 | 5.00 | **0.028** |
|  | Climate scenario × aphid infestation × plant composition | 2,84 | 0.71 | 0.495 |
| Total proteins | Climate scenario | 2,84 | 23.41 | **<0.001** |
|  | Plant composition | 1,84 | 1.42 | 0.238 |
|  | Aphid infestation | 1,84 | 30.03 | **<0.001** |
|  | Climate scenario × aphid infestation | 2,84 | 1.36 | 0.262 |
|  | Climate scenario × plant composition | 2,84 | 2.81 | 0.066 |
|  | Plant composition × aphid infestation | 1,84 | 0.01 | 0.924 |
|  | Climate scenario × aphid infestation × plant composition | 2,84 | 2.46 | **0.025** |
| Lipids | Climate scenario | 2.84 | 29.66 | **<0.001** |
|  | Plant composition | 1,84 | 13.88 | **<0.001** |
|  | Aphid infestation | 1.84 | 61.04 | **<0.001** |
|  | Climate scenario × aphid infestation | 2.84 | 5.93 | **0.004** |
|  | Climate scenario × plant composition | 2,84 | 0.86 | 0.426 |
|  | Plant composition × aphid infestation | 1,84 | 1.21 | 0.274 |
|  | Climate scenario × aphid infestation × plant composition | 2,84 | 7.38 | **0.001** |

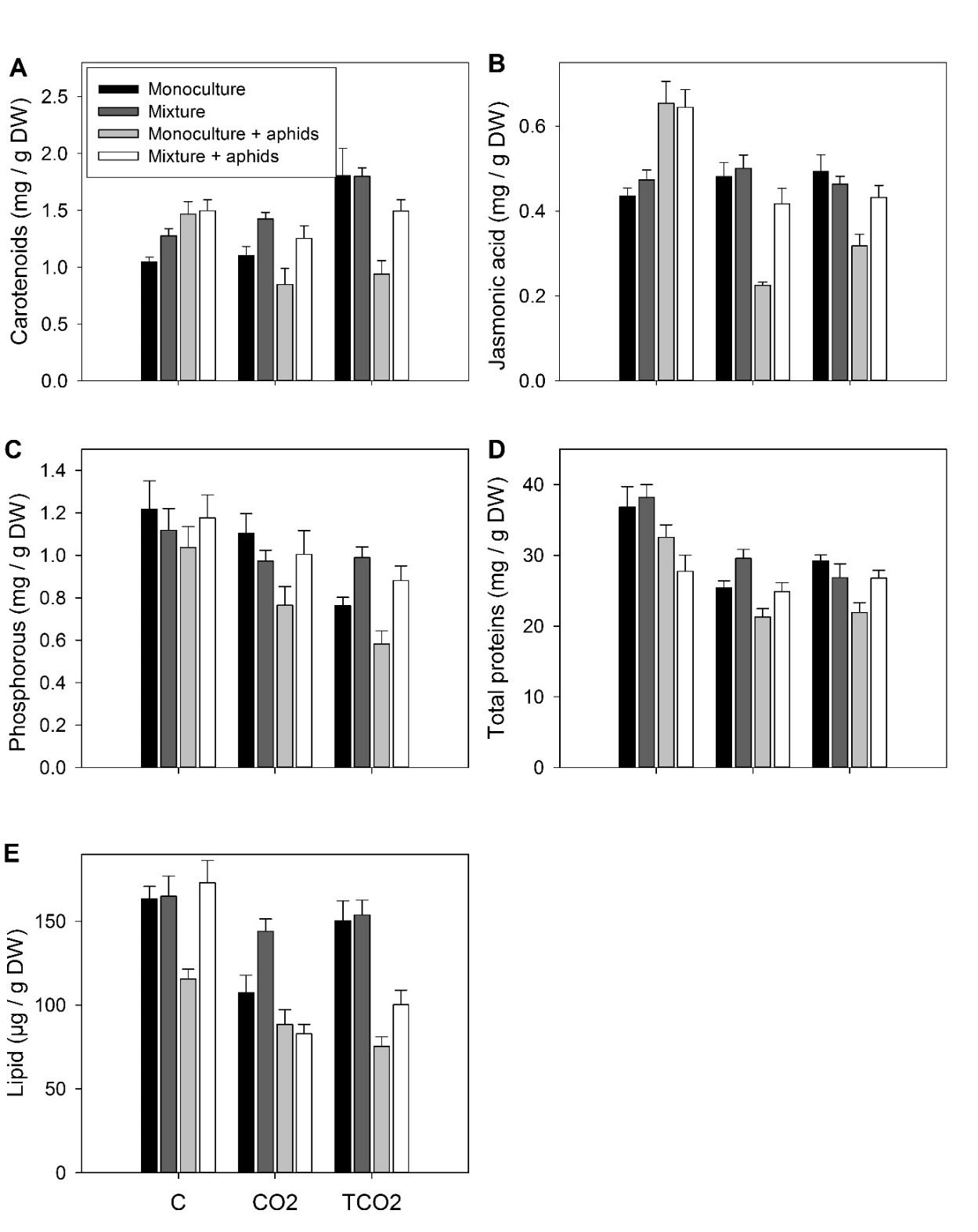
**Fig. S3.2** Effect of climate scenario (C, CO_2_ and TCO_2_), aphid infestation and plant composition on the metabolite levels in cluster 2. Bars represent means ± SE. Plant communities consist of monocultures of *P. lanceolata* and mixtures of *Lolium perenne* and *P. lanceolata*.

**Table S3.3** Summary of GLM results for effects of climate scenario, plant composition and aphid infestation on the metabolite levels in cluster 3. Plant communities consist of monocultures of *P. lanceolata* and mixtures of *Lolium perenne* and *P. lanceolata*. P-values are presented in bold when significant (≤ 0.05).

| Measurement | Treatment | df | F | P |
| --- | --- | --- | --- | --- |
| Leaf nitrogen | Climate scenario | 2,9 | 2.89 | 0.108 |
|  | Plant composition | 1,75 | 9.82 | **0.003** |
|  | Aphid infestation | 1,75 | 4.74 | **0.033** |
|  | Climate scenario × aphid infestation | 2,75 | 0.25 | 0.780 |
|  | Climate scenario × plant composition | 2,75 | 0.49 | 0.615 |
|  | Plant composition × aphid infestation | 1,75 | 1.95 | 0.167 |
|  | Climate scenario × aphid infestation × plant composition | 2,75 | 0.15 | 0.857 |

**Table S3.4** Summary of GLM results for effects of climate scenario, plant composition and aphid infestation on the metabolite levels in cluster 4. Plant communities consist of monocultures of *P. lanceolata* and mixtures of *Lolium perenne* and *P. lanceolata*. P-values are presented in bold when significant (≤ 0.05).

| Measurement | Treatment | df | F | P |
| --- | --- | --- | --- | --- |
| Leaf carbon | Climate scenario | 2,84 | 3.00 | 0.055 |
|  | Plant composition | 1,84 | 8.00 | **0.006** |
|  | Aphid infestation | 1,84 | 0.00 | 0.945 |
|  | Climate scenario × aphid infestation | 2,84 | 1.65 | 0.199 |
|  | Climate scenario × plant composition | 2,84 | 0.16 | 0.852 |
|  | Plant composition × aphid infestation | 1,84 | 1.37 | 0.245 |
|  | Climate scenario × aphid infestation × plant composition | 2,84 | 0.41 | 0.667 |
| Cellulose | Climate scenario | 2,84 | 0.14 | 0.870 |
|  | Plant composition | 1,84 | 6.99 | **0.010** |
|  | Aphid infestation | 1,84 | 8.66 | **0.004** |
|  | Climate scenario × aphid infestation | 2,84 | 0.90 | 0.409 |
|  | Climate scenario × plant composition | 2,84 | 0.14 | 0.866 |
|  | Plant composition × aphid infestation | 1,84 | 12.04 | **0.001** |
|  | Climate scenario × aphid infestation × plant composition | 2,84 | 4.05 | **0.021** |
| C:P ratio | Climate scenario | 2,84 | 15.98 | **<0.001** |
|  | Plant composition | 1,84 | 10.04 | **0.002** |
|  | Aphid infestation | 1,84 | 8.66 | **0.004** |
|  | Climate scenario × aphid infestation | 2,84 | 0.80 | 0.452 |
|  | Climate scenario × plant composition | 2,84 | 3.66 | **0.030** |
|  | Plant composition × aphid infestation | 1,84 | 5.61 | **0.020** |
|  | Climate scenario × aphid infestation × plant composition | 2,84 | 0.35 | 0.703 |
| N:P ratio | Climate scenario | 2,9 | 7.81 | **0.011** |
|  | Plant composition | 1,75 | 23.32 | **<0.001** |
|  | Aphid infestation | 1,75 | 4.39 | **0.040** |
|  | Climate scenario × aphid infestation | 2,75 | 1.50 | 0.230 |
|  | Climate scenario × plant composition | 2,75 | 7.60 | **0.001** |
|  | Plant composition × aphid infestation | 1,75 | 3.82 | 0.054 |
|  | Climate scenario × aphid infestation × plant composition | 2,75 | 0.20 | 0.820 |
| Total antioxidant capacity | Climate scenario | 2,84 | 5.97 | **0.004** |
|  | Plant composition | 1,84 | 0.80 | 0.374 |
|  | Aphid infestation | 1,84 | 261.74 | **<0.001** |
|  | Climate scenario × aphid infestation | 2,84 | 0.12 | 0.889 |
|  | Climate scenario × plant composition | 2,84 | 4.00 | **0.022** |
|  | Plant composition × aphid infestation | 1,84 | 7.31 | **0.008** |
|  | Climate scenario × aphid infestation × plant composition | 2,84 | 0.15 | 0.865 |
| Catalpol | Climate scenario | 2,84 | 1.38 | 0.257 |
|  | Plant composition | 1,84 | 6.73 | **0.011** |
|  | Aphid infestation | 1,84 | 152.81 | **<0.001** |
|  | Climate scenario × aphid infestation | 2,84 | 0.99 | 0.375 |
|  | Climate scenario × plant composition | 2,84 | 0.74 | 0.480 |
|  | Plant composition × aphid infestation | 1,84 | 5.92 | **0.017** |
|  | Climate scenario × aphid infestation × plant composition | 2,84 | 0.20 | 0.821 |
| Aucubin | Climate scenario | 2,84 | 5.53 | **0.005** |
|  | Plant composition | 1,84 | 1.31 | 0.255 |
|  | Aphid infestation | 1,84 | 128.12 | **<0.001** |
|  | Climate scenario × aphid infestation | 2,84 | 5.47 | **0.006** |
|  | Climate scenario × plant composition | 2,84 | 1.29 | 0.280 |
|  | Plant composition × aphid infestation | 1,84 | 1.31 | 0.255 |
|  | Climate scenario × aphid infestation × plant composition | 2,84 | 1.04 | 0.357 |
| Sucrose | Climate scenario | 2,84 | 9.24 | **<0.001** |
|  | Plant composition | 1,84 | 21.14 | **<0.001** |
|  | Aphid infestation | 1,84 | 30.22 | **<0.001** |
|  | Climate scenario × aphid infestation | 2,84 | 8.07 | **0.001** |
|  | Climate scenario × plant composition | 2,84 | 8.21 | **0.001** |
|  | Plant composition × aphid infestation | 1,84 | 23.89 | **<0.001** |
|  | Climate scenario × aphid infestation × plant composition | 2,84 | 2.52 | 0.0863 |
| Leaf membrane damage, MDA | Climate scenario | 2,84 | 19.04 | **<0.001** |
|  | Plant composition | 1,84 | 34.17 | **<0.001** |
|  | Aphid infestation | 1,84 | 31.02 | **<0.001** |
|  | Climate scenario × aphid infestation | 2,84 | 0.07 | 0.935 |
|  | Climate scenario × plant composition | 2,84 | 2.65 | 0.077 |
|  | Plant composition × aphid infestation | 1,84 | 3.84 | 0.054 |
|  | Climate scenario × aphid infestation × plant composition | 2,84 | 2.31 | 0.105 |
| Proline | Climate scenario | 2,84 | 29.37 | **<0.001** |
|  | Plant composition | 1,84 | 22.55 | **<0.001** |
|  | Aphid infestation | 1,84 | 138.62 | **<0.001** |
|  | Climate scenario × aphid infestation | 2,84 | 24.19 | **<0.001** |
|  | Climate scenario × plant composition | 2,84 | 11.63 | **<0.001** |
|  | Plant composition × aphid infestation | 1,84 | 18.05 | **<0.001** |
|  | Climate scenario × aphid infestation × plant composition | 2,84 | 4.78 | **0.011** |
| Salicylic acid | Climate scenario | 2,84 | 15.71 | **<0.001** |
|  | Plant composition | 1,84 | 150.51 | **<0.001** |
|  | Aphid infestation | 1,84 | 390.48 | **<0.001** |
|  | Climate scenario × aphid infestation | 2,84 | 21.69 | **<0.001** |
|  | Climate scenario × plant composition | 2,84 | 6.70 | **0.002** |
|  | Plant composition × aphid infestation | 1,84 | 110.21 | **<0.001** |
|  | Climate scenario × aphid infestation × plant composition | 2,84 | 38.79 | **<0.001** |
| Lignin | Climate scenario | 2,9 | 16.45 | **0.001** |
|  | Plant composition | 1,75 | 67.05 | **<0.001** |
|  | Aphid infestation | 1,75 | 81.49 | **<0.001** |
|  | Climate scenario × aphid infestation | 2,75 | 14.64 | **<0.001** |
|  | Climate scenario × plant composition | 2,75 | 8.24 | **0.001** |
|  | Plant composition × aphid infestation | 1,75 | 8.27 | **0.005** |
|  | Climate scenario × aphid infestation × plant composition | 2,75 | 3.14 | **0.049** |
| Lignin:N | Climate scenario | 2,9 | 24.88 | **<0.001** |
|  | Plant composition | 1,75 | 22.36 | **<0.001** |
|  | Aphid infestation | 1,75 | 63.02 | **<0.001** |
|  | Climate scenario × aphid infestation | 2,75 | 2.26 | 0.112 |
|  | Climate scenario × plant composition | 2,75 | 3.43 | **0.037** |
|  | Plant composition × aphid infestation | 1,75 | 5.18 | **0.026** |
|  | Climate scenario × aphid infestation × plant composition | 2,75 | 1.07 | 0.348 |

**Fig. S3.3** Effect of climate scenario (C, CO_2_ and TCO_2_), aphid infestation and plant composition on the metabolite levels (sugars, C:N, N:P, TAC; Aucubin, Catalpol & cellulose) in cluster 4. Bars represent means ± SE. Plant communities consist of monocultures of *P. lanceolata* and mixtures of *Lolium perenne* and *P. lanceolata*.

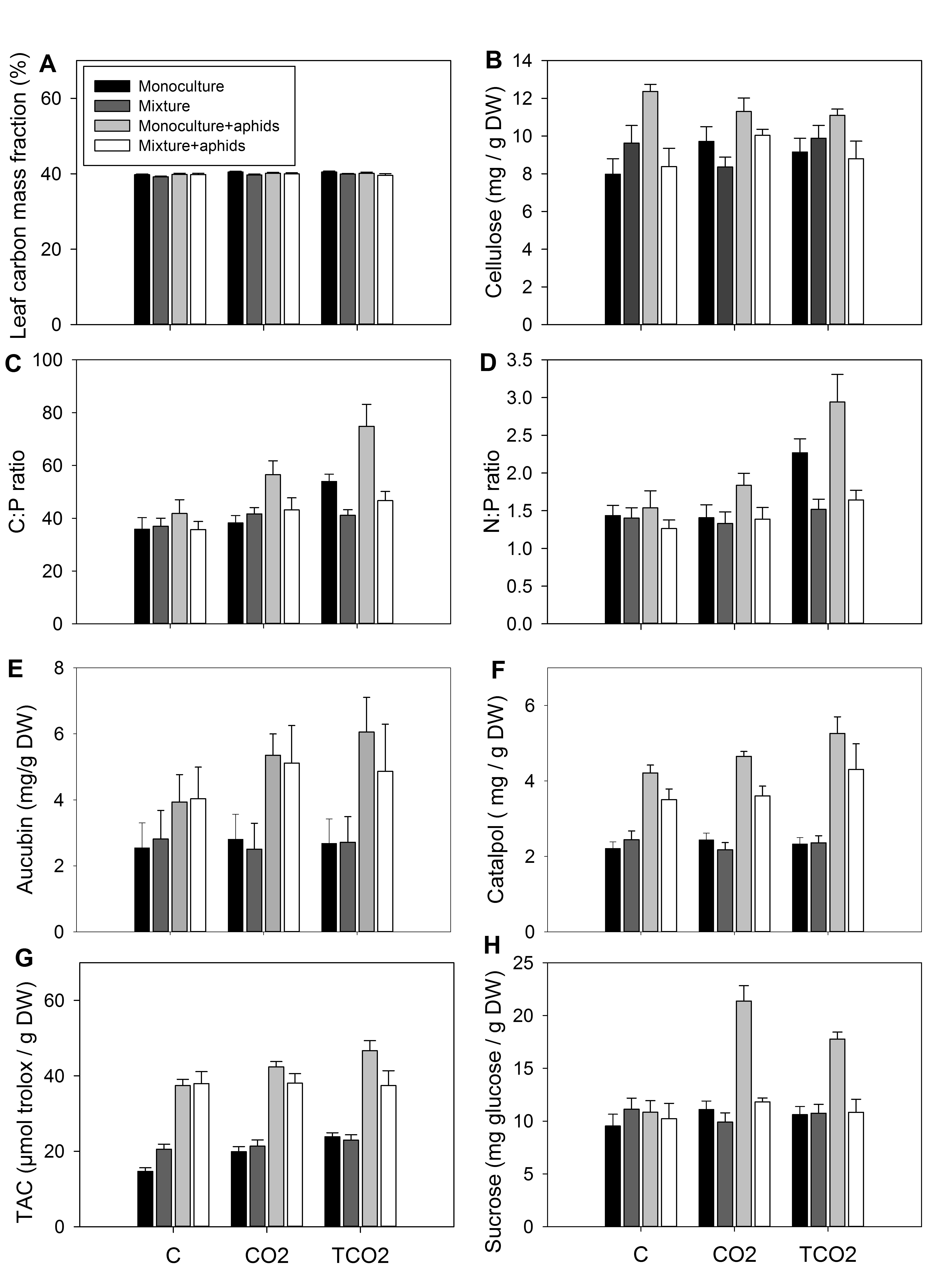

**Fig. S3.4** Effect of climate scenario (C, CO_2_ and TCO_2_), aphid infestation and plant composition on the on the metabolite levels (MDA; lignin, salicylic acid & proline) in cluster 4. Bars represent means ± SE. Plant communities consist of monocultures of *P. lanceolata* and mixtures of *Lolium perenne* and *P. lanceolata*.

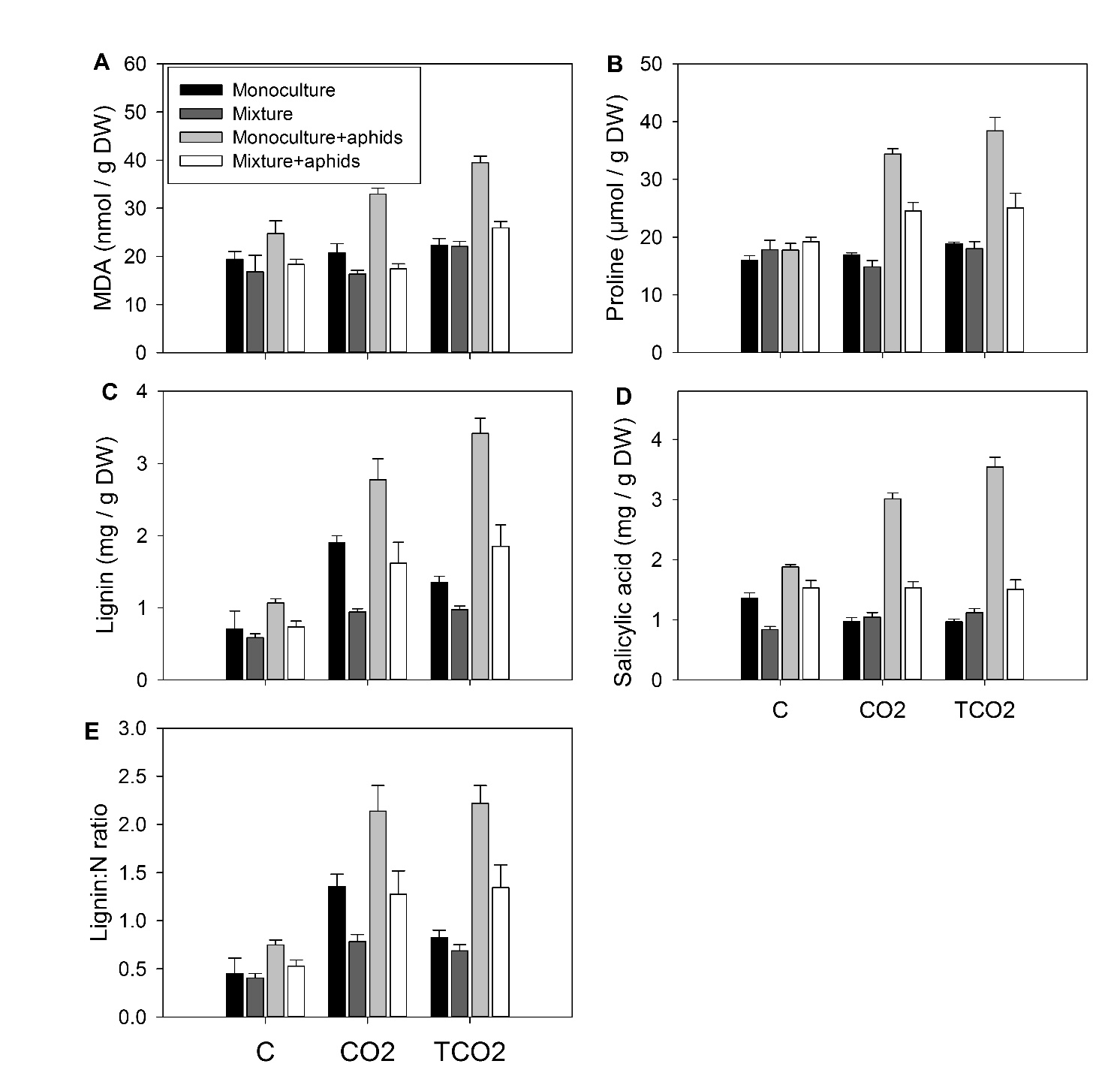
