## Appendix 1 for "Interspecific plant competition mediates the metabolic and ecological signature of a plant-herbivore interaction under warming and elevated CO2"

**Appendix 1: schematic representation of the setup and clustering of the metabolites without total soluble sugars**

**Fig S1.1:** Schematic representation of the experimental setup

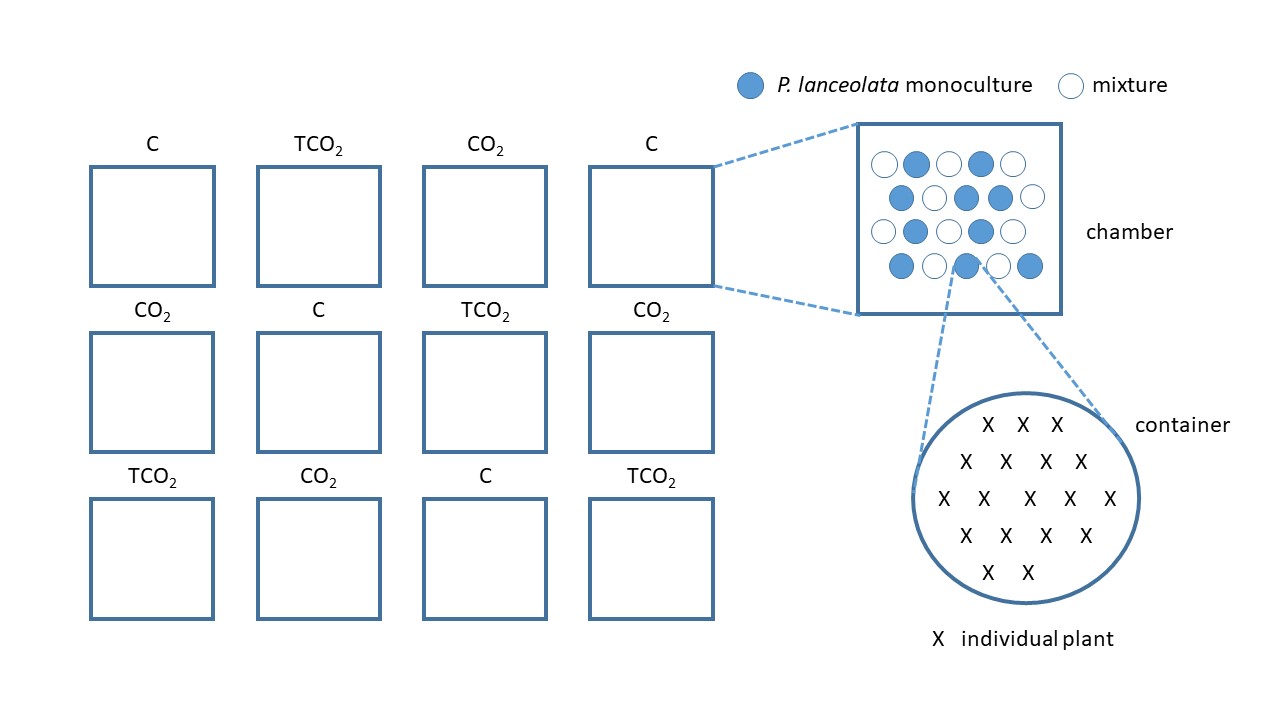

**Fig S1.2:** Clustering with removing total amount of soluble sugars

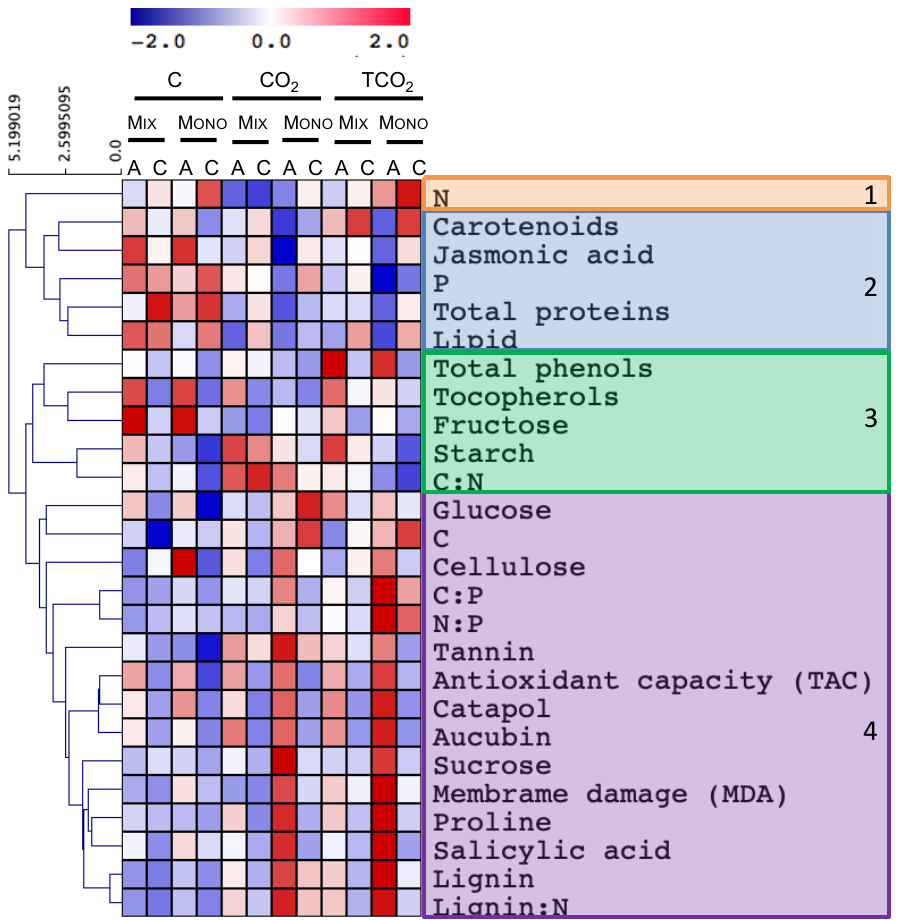
