## Appendix 2 for "Interspecific plant competition mediates the metabolic and ecological signature of a plant-herbivore interaction under warming and elevated CO2"

**Supporting information 2: *Extraction procedure and SEM statistics***

***Extraction procedure***

*Carbohydrates*

Small soluble sugars were determined in 0.2 g (DW) plant material, ground in liquid nitrogen (MagNALyser, Roche, Vilvoorde, Belgium) and extracted in 1 ml of 50 mM TAE buffer pH 7.5 (0.02% sodium azide, 10 mM mannitol, 0.1% polyclar, 10 mM NaHSO_3_, 1 mM mercapto-ethanol, 1 mM phenylmethanesulfonylfluoride (PMSF)). The extract was centrifuged (14,000 g, 4 °C, 5 min), 150 µl was heated for 5 min in a water bath at 90 °C. After cooling and centrifugation (14,000 g, 4 °C, 5 min), the supernatant was added to a mixed bed Dowex column (300 µl Dowex H+, 300 µl Dowex Ac–; both 100–200 mesh; Acros Organics, Morris Plains, NJ, USA). The column was eluted six times with 150 µl of ddH2O. Glucose, fructose and sucrose concentrations were measured by HPAEC-PAD as before (Vergauwen *et al.*, 2000). Total soluble sugars and starch content were estimated by the anthrone reagent method (Leyva *et al.*, 2008).

*Total protein*

A plant sample (200 mg DW) was homogenized in 2 ml of cold 0.05M K-Phosphate buffer (pH7.0) and centrifuged at 15,000 g at 4 °C for 20 min. The supernatant was treated by 10% (w/v) TCA to precipitate soluble protein, which was redissolved in 1 N NaOH. The remaining pellet was used to extract insoluble protein. It was successively washed with 80% ethanol, 10% (w/v) cold TCA, ethanol:chloroform (3:1, v/v), ethanol:ether (3:1, v/v), and ether to remove phenolic compounds. The washed pellet was then dissolved in 1 N NaOH at 80 °C for 1 h. Soluble and insoluble protein content was estimated according to Lowry *et al.* (1951). Total protein content was calculated by adding the contents of soluble and insoluble proteins.

*Lignin, polyphenols and tannin*

For lignin determination, MagNALyser homogenized 0.1 g DW with 95% ethanol. The homogenate was centrifuged at 14 g for 3 min. Successively, the pellet was washed with 95% ethanol 30 min at 76 °C, chloroform for 30 min at 59 °C and then incubated in acetone for 30 min at 54 °C. One ml of 25% acetyl bromide in acetic acid (1:3, v/v) was added to the pellet and incubated at 70 °C for 30 min. After cooling, 0.2 ml of 2 M NaOH and 0.1 ml of 7.5 M hydroxylamine hydrochloride were added, and the volume was made up to 10 ml with acetic acid. After centrifugation at 1000 g for 5 min, the absorbance of the supernatant was measured against a NaOH blank at 280 nm (Lin & Kao, 2001). Polyphenol contents were extracted in 80% ethanol (v/v) and determined according to Zhang *et al.* (2006), with gallic acid as standard. Tannin content was determined as described by Hagerman and Butler (1978). About 0.2 g FW was homogenized in 2 ml acetate buffer pH 5 containing 2 mg of bovine serum albumin. The mixture was incubated for 15 min at room temperature and then centrifuged at 14,000 g for 15 min then the pellet was dissolved in 4 ml of a solution consisting of 1% SDS and 5% of tri-ethanolamine in water. One ml of 10 mM FeCl3 in 0.01 N HCl was added and incubated for 15 min. Then the absorbance was determined at 510 nm. Tannic acid was used as the standard.

*Membrane damage (lipid peroxidation (MDA))*

Lipid peroxidation was determined on 200 mg dry tissues, homogenized in 2 ml 80% ethanol by mortar and pestle, using a thiobarbituric acid-malondialdehyde (TBA-MDA) assay (Hodges *et al.*, 1999).

*Total antioxidant capacity*

Plant tissues (200 mg DW) were ground by a MagNALyser in liquid nitrogen and the antioxidants were extracted in 2 ml of ice cold 80% ethanol. FRAP (ferric reducing/antioxidant power assay) reagent (0.3 M acetate buffer (pH3.6), 0.01 mM TPTZ in 0.04 mM HCl, 0.02 M FeCl3.6H2O) was mixed with the extract and measured at 600 nm using a microplate reader (Synergy Mx, Biotek Instruments Inc., Vermont, USA) (Benzie & Strain, 1999). Trolox was used as standard.

*Tocopherols*

Tocopherols were extracted with hexane using the MagNALyser. The dried extract (CentriVap concentrator, Labconco, Kansas, USA) was resuspended in hexane, and tocopherols were separated and quantified by HPLC (Shimadzu, ‘s Hertogenbosch, The Netherlands) (normal phase conditions, Particil Pac 5 µm column material, length 250 mm, i.d. 4.6 mm). Dimethyl tocol (DMT) was used as internal standard (5 ppm). Data were analysed with Shimadzu Class VP 6.14 software.

*Catalpol and aucubin*

Each sample was extracted overnight in 70 % methanol, and then filtered (12–15 μm) followed by a dilution of 10 times with ultrapure water. The concentrations of the aucubin and catalpol were analyzed using HPLC as described by Marak *et al.* (2002).

*Salicylic acid and jasmonic acid*

Salicylic acid concentration was measured according Li *et al.* (1999). This procedure had a 25% recovery rate, as determined by extracting known amounts of salicylic acid. Samples were homogenized in the extraction buffer. The samples were analysed by GC-MS after addition of 150 ng of 13C1,2-JA as an internal standard (Schittko *et al.*, 2000). The tissue was homogenized with a reciprocating shaker at 6.0 m sec ± 1 for 90 sec in extraction tubes containing 900 mg of lysing matrix (BIO 101, Vista, California, USA).

***SEM statistics***

**Table S2.1** Partial slopes of the structural equation model presented in Figure 2A. Labels CO_2_ and TCO_2_ indicate elevated CO_2_ and combined warming and elevated CO_2_, respectively. Plant communities consist of monocultures of *Plantago lanceolata* and mixtures of *P. lanceolata* and *Lolium perenne*. The four clusters refer to those obtained by the hierarchical clustering analysis (see Fig. 1). P values are presented in bold when significant (≤0.05).

| Response | Predictor | Estimate | SE | P-value |
| --- | --- | --- | --- | --- |
| Mildew | TCO_2_ | 0.563 | 0.232 | **0.038** |
|  | CO_2_ | -0.125 | 0.232 | 0.603 |
| Cluster 1 | Aphid infestation | 1.171 | 0.217 | **0.000** |
|  | CO_2_ | 1.024 | 0.224 | **0.001** |
|  | CO_2_ × aphid infestation | -0.899 | 0.306 | **0.007** |
|  | Plant composition | 0.511 | 0.219 | **0.028** |
|  | CO_2_ × plant composition | -0.586 | 0.308 | 0.068 |
|  | TCO_2_ | 0.438 | 0.233 | 0.093 |
|  | TCO_2_ × aphid infestation | -0.324 | 0.306 | 0.300 |
|  | Plant composition × aphid infestation | -0.264 | 0.308 | 0.399 |
|  | TCO_2_ × plant composition × aphid infestation | 0.330 | 0.434 | 0.454 |
|  | CO_2_ × plant composition × aphid infestation | 0.329 | 0.433 | 0.454 |
|  | TCO_2_ × plant composition | -0.163 | 0.308 | 0.600 |
|  | Mildew | 0.036 | 0.129 | 0.780 |
| Cluster 2 | TCO_2_ × aphid infestation | -1.382 | 0.323 | **0.000** |
|  | CO_2_ × aphid infestation | -0.952 | 0.323 | **0.007** |
|  | CO_2_ | -0.591 | 0.229 | **0.030** |
|  | TCO_2_ × plant composition × aphid infestation | 0.746 | 0.459 | 0.116 |
|  | CO_2_ × plant composition | 0.244 | 0.325 | 0.459 |
|  | TCO_2_ | -0.151 | 0.237 | 0.541 |
|  | Plant composition | 0.138 | 0.231 | 0.555 |
|  | CO_2_ × plant composition × aphid infestation | 0.228 | 0.457 | 0.622 |
|  | Plant composition × aphid infestation | 0.090 | 0.325 | 0.785 |
|  | TCO_2_ × plant composition | -0.086 | 0.325 | 0.793 |
|  | Aphid infestation | 0.028 | 0.229 | 0.905 |
|  | Mildew | -0.010 | 0.127 | 0.939 |
| Cluster 3 | Aphid infestation | -0.748 | 0.397 | 0.071 |
|  | Plant composition | -0.549 | 0.403 | 0.185 |
|  | CO_2_ | -0.646 | 0.483 | 0.213 |
|  | Plant composition × aphid infestation | 0.432 | 0.566 | 0.453 |
|  | Mildew | 0.198 | 0.282 | 0.489 |
|  | TCO_2_ × plant composition | -0.346 | 0.566 | 0.547 |
|  | TCO_2_ × aphid infestation | 0.214 | 0.561 | 0.707 |
|  | CO_2_ × plant composition | -0.210 | 0.566 | 0.714 |
|  | CO_2_ × aphid infestation | 0.174 | 0.561 | 0.759 |
|  | CO_2_ × plant composition × aphid infestation | 0.239 | 0.794 | 0.766 |
|  | TCO_2_ | 0.151 | 0.503 | 0.771 |
|  | TCO_2_ × plant composition × aphid infestation | -0.182 | 0.797 | 0.821 |
| Cluster 4 | Aphid infestation | 0.729 | 0.154 | 0.000 |
|  | TCO_2_ × aphid infestation | 0.987 | 0.218 | 0.000 |
|  | CO2 × aphid infestation | 0.631 | 0.218 | 0.008 |
|  | TCO_2_ | 0.525 | 0.160 | 0.010 |
|  | TCO_2_ × plant composition × aphid infestation | -0.766 | 0.309 | 0.020 |
|  | CO_2_ | 0.401 | 0.154 | 0.029 |
|  | CO_2_ × plant composition | -0.388 | 0.219 | 0.089 |
|  | Plant composition × aphid infestation | -0.385 | 0.219 | 0.091 |
|  | TCO_2_ × plant composition | -0.227 | 0.219 | 0.310 |
|  | CO_2_ × plant composition × aphid infestation | -0.251 | 0.309 | 0.423 |
|  | Mildew | -0.062 | 0.086 | 0.472 |
|  | Plant composition | 0.011 | 0.156 | 0.944 |
| Biomass *P. lanceolata* | CO_2_ × plant composition × aphid infestation | 1.088 | 0.571 | 0.068 |
|  | TCO_2_ | -0.904 | 0.488 | 0.097 |
|  | CO_2_ × aphid infestation | -0.626 | 0.404 | 0.133 |
|  | CO_2_ | 0.566 | 0.474 | 0.263 |
|  | Mildew | 0.142 | 0.229 | 0.542 |
|  | TCO_2_ × plant composition | -0.203 | 0.408 | 0.622 |
|  | Aphid infestation | -0.128 | 0.286 | 0.659 |
|  | TCO_2_ × plant composition × aphid infestation | 0.186 | 0.574 | 0.748 |
|  | CO_2_ × plant composition | -0.113 | 0.408 | 0.784 |
|  | Plant composition | 0.060 | 0.291 | 0.838 |
|  | TCO_2_ × aphid infestation | -0.064 | 0.404 | 0.876 |
|  | Plant composition × aphid infestation | -0.021 | 0.408 | 0.960 |

**Table S2.2** Partial slopes of the structural equation model presented in Figure 2B. Labels CO_2_ and TCO_2_ indicate elevated CO_2_ and combined warming and elevated CO_2_, respectively. Plant communities consist of monocultures of *Plantago lanceolata* and mixtures of *P. lanceolata* and *Lolium perenne*. The four clusters refer to those obtained by the hierarchical clustering analysis (see Fig. 1). P values are presented in bold when significant (≤0.05).

| Response | Predictor | Estimate | SE | P-value |
| --- | --- | --- | --- | --- |
| Aphid population | Cluster 2 control | 1.558 | 0.664 | 0.066 |
|  | CO_2_ | 1.650 | 1.462 | 0.288 |
|  | TCO_2_ | 1.149 | 1.088 | 0.319 |
|  | Cluster 1 control | -0.837 | 0.990 | 0.436 |
|  | Plant composition | -0.463 | 0.581 | 0.461 |
|  | Cluster 3 control | -0.218 | 0.310 | 0.512 |
|  | TCO_2_ × plant composition | 0.210 | 0.653 | 0.761 |
|  | CO_2_ × plant composition | -0.315 | 1.043 | 0.774 |
|  | Cluster 4 control | 0.185 | 1.140 | 0.878 |
| Cluster 1 control | CO_2_ | 1.024 | 0.150 | **0.000** |
|  | Plant composition | 0.502 | 0.131 | **0.004** |
|  | CO_2_ × plant composition | -0.595 | 0.185 | **0.011** |
|  | TCO_2_ | 0.456 | 0.150 | **0.014** |
|  | TCO_2_ × plant composition | -0.154 | 0.185 | 0.426 |
| Cluster 2 control | CO_2_ | -0.591 | 0.235 | **0.033** |
|  | CO_2_ × plant composition | 0.247 | 0.241 | 0.333 |
|  | Plant composition | 0.140 | 0.171 | 0.431 |
|  | TCO_2_ | -0.156 | 0.235 | 0.525 |
|  | TCO_2_ × plant composition | -0.089 | 0.241 | 0.722 |
| Cluster 3 control | Plant composition | -0.599 | 0.339 | 0.111 |
|  | CO_2_ | -0.646 | 0.524 | 0.248 |
|  | TCO_2_ × plant composition | -0.296 | 0.479 | 0.552 |
|  | CO_2_ × plant composition | -0.259 | 0.479 | 0.602 |
|  | TCO_2_ | 0.250 | 0.524 | 0.645 |
| Cluster 4 control | TCO_2_ | 0.494 | 0.115 | **0.002** |
|  | CO_2_ | 0.401 | 0.115 | **0.007** |
|  | CO_2_ × plant composition | -0.372 | 0.148 | **0.033** |
|  | TCO_2_ × plant composition | -0.243 | 0.148 | 0.136 |
|  | Plant composition | 0.027 | 0.105 | 0.806 |
| Cluster 1 aphids | Plant composition | 0.518 | 0.327 | 0.212 |
|  | TCO_2_ × aphid population × plant composition | -0.675 | 0.471 | 0.247 |
|  | CO_2_ × plant composition | -0.526 | 0.402 | 0.281 |
|  | CO_2_ × aphid population × plant composition | -0.536 | 0.462 | 0.329 |
|  | CO_2_ × aphid population | -0.269 | 0.281 | 0.409 |
|  | Aphid population × plant composition | 0.329 | 0.398 | 0.469 |
|  | Aphid population | 0.133 | 0.200 | 0.555 |
|  | CO_2_ | 0.115 | 0.257 | 0.664 |
|  | TCO_2_ | 0.050 | 0.268 | 0.856 |
|  | TCO_2_ × aphid population | 0.017 | 0.255 | 0.952 |
|  | TCO_2_ × plant composition | 0.002 | 0.410 | 0.997 |
| Cluster 2 aphids | CO_2_ | -1.525 | 0.200 | **0.000** |
|  | TCO_2_ | -1.590 | 0.208 | **0.000** |
|  | TCO_2_ × aphid population × plant composition | -0.892 | 0.394 | 0.108 |
|  | CO_2_ × aphid population × plant composition | -0.780 | 0.388 | 0.138 |
|  | TCO_2_ × aphid population | 0.375 | 0.199 | 0.155 |
|  | TCO_2_ × plant composition | 0.564 | 0.341 | 0.197 |
|  | Plant composition | 0.431 | 0.269 | 0.207 |
|  | Aphid population × plant composition | 0.531 | 0.333 | 0.209 |
|  | Aphid population | -0.227 | 0.155 | 0.241 |
|  | CO_2_ × aphid population | 0.266 | 0.218 | 0.310 |
|  | CO_2_ × plant composition | 0.278 | 0.334 | 0.466 |
| Cluster 3 aphids | CO_2_ × aphid population × plant composition | 0.993 | 0.489 | 0.135 |
|  | Aphid population × plant composition | -0.857 | 0.440 | 0.147 |
|  | TCO_2_ × aphid population × plant composition | 0.675 | 0.500 | 0.270 |
|  | TCO_2_ | 0.508 | 0.470 | 0.309 |
|  | CO_2_ | -0.492 | 0.460 | 0.312 |
|  | Plant composition | -0.501 | 0.429 | 0.327 |
|  | TCO_2_ × aphid population | -0.400 | 0.402 | 0.392 |
|  | CO_2_ × aphid population | -0.433 | 0.481 | 0.434 |
|  | Aphid population | 0.264 | 0.344 | 0.498 |
|  | CO_2_ × plant composition | 0.363 | 0.476 | 0.501 |
|  | TCO_2_ × plant composition | -0.156 | 0.483 | 0.768 |
| Cluster 4 aphids | TCO_2_ | 1.432 | 0.207 | **0.000** |
|  | CO_2_ | 1.015 | 0.199 | **0.001** |
|  | TCO_2_ × plant composition | -0.954 | 0.339 | 0.067 |
|  | CO_2_ × plant composition | -0.618 | 0.333 | 0.160 |
|  | CO_2_ × aphid population | -0.337 | 0.217 | 0.219 |
|  | Plant composition | -0.377 | 0.267 | 0.253 |
|  | Aphid population | 0.214 | 0.155 | 0.260 |
|  | TCO_2_ × aphid population | -0.150 | 0.198 | 0.504 |
|  | Aphid population × plant composition | -0.191 | 0.331 | 0.605 |
|  | CO_2_ × aphid population × plant composition | 0.218 | 0.386 | 0.611 |
|  | TCO_2_ × aphid population × plant composition | 0.096 | 0.391 | 0.823 |
